# TDG orchestrates ATF4-dependent gene transcription during retinoic acid-induced cell fate acquisition

**DOI:** 10.1101/2024.04.01.587571

**Authors:** Marion Turpin, Thierry Madigou, Maud Bizot, Rachael Acker, Stephane Avner, Gérard Benoît, Martin Braud, Cynthia Fourgeux, Gaëlle Palierne, Jeremie Poschmann, Katie Sawvell, Erwan Watrin, Christine Le Péron, Gilles Salbert

## Abstract

Acquisition of cell identity is associated with a remodeling of the epigenome in part through active DNA demethylation. The T:G mismatch DNA glycosylase (TDG) participates to this process by removing 5-methylcytosines that have been oxidized by Ten-Eleven-Translocation (TET) enzymes. Despite this well-defined molecular function, a comprehensive view of the biological function of TDG is still lacking, especially during cell differentiation. Here, we combined transcriptomic and epigenomic approaches in a *Tdg* knock-out epiblast stem-like cell model to decipher TDG function in pluripotent cells and their retinoic acid-induced progeny. We determined that TDG occupies a majority of active promoters, a large fraction of which are also engaged by the transcription factor ATF4. Consistently, neural fate commitment upon retinoic acid treatment is associated with a TDG-dependent sustained expression of ATF4-dependent genes, in relation with a TDG-associated nucleosome positioning at promoters. We further evidenced that TDG maintains ATF4 pathway activity by positively regulating the mammalian target of rapamycin complex 1 (mTORC1), favoring neural cell fate commitment. These observations highlight the central role of TDG in cell differentiation and support a model linking metabolic reprogramming to cell fate acquisition.

## INTRODUCTION

During differentiation, acquisition of a given cell identity relies on the establishment of specific gene expression profiles. These transcriptional changes involve both remodeling and covalent modifications of chromatin, among which stands DNA methylation. Genome-wide DNA methylation patterns constitute a crucial layer of epigenetic information involved in transcription regulation and result from the equilibrium between methylation reactions catalyzed by DNA methyltransferases (DNMTs) and active demethylation reactions, which rely on the Ten-Eleven-Translocation (TET) and thymidine:guanidine mismatch thymine DNA glycosylase (TDG) enzymes (Wu and Zhang, 2017). The transfer by DNMTs of a methyl group from S-adenosyl-L-methionine to cytosine at position 5 (5mC) occurs mainly in the context of CpG dinucleotides and can exert either negative or positive effects on gene expression by regulating the binding of transcription factors to DNA, and by the interaction of 5mCpGs with methyl-DNA binding proteins which are believed to recruit repressive complexes (Nan et al., 1998; Kaluscha et al., 2022). However, in the context of CpG island (CGIs) promoters, dense DNA methylation is associated with gene silencing (Deaton and Bird, 2011).

TDG was initially discovered as an enzyme catalysing the removal of U and T mismatched with G (Neddermann and Jiricny, 1993; 1994) thereby participating in the maintenance of genome stability. Its role has expanded since the discovery of an active DNA demethylation pathway that involves TDG and TET enzymes (Kriaucionis and Heintz, 2009; Tahiliani et al., 2009; Maïti and Drohat, 2011). During this demethylation process, TETs iteratively oxidize 5mC into 5-hydroxy-mC (5hmC), 5-formyl-C (5fC) and 5-carboxyl-C (5caC), collectively referred to as oxi-mCs. The two terminal oxidized forms, 5fC and 5caC, are excised by TDG that hydrolyses the sugar-base bond, creating an abasic site that is then converted into an intact, unmodified cytosine by the base excision repair machinery. The newly incorporated base can then be targeted for *de novo* methylation by DNA methyltransferases. One crucial aspect of DNA methylation resides in the dynamic state that exists between modified and unmodified forms, which varies among different types of genomic regions. The turnover of 5mC is notably high at enhancers, a particular class of gene regulatory elements that physically contact distant promoters for expression of cognate genes. As enhancer/promoter wiring plays an essential role in cell specific gene expression patterns, it is tempting to consider that such highly dynamic methylation of CpGs at enhancers is indicative of its involvement in programming and acquisition of cell identity (Sérandour et al., 2012; Caron et al., 2015; Mahé et al., 2017; Ginno et al., 2020). As an actor of the demethylation pathway, TDG has been involved in this cyclic methylation/demethylation process in breast cancer cells, a role that was later extended to embryonic stem cells (ESCs - Métivier et al., 2008; Rulands et al. 2018; Parry et al., 2021). Furthermore, TDG interaction with DNMT3A and DNMT3B results in a decreased catalytic activity of both methyltransferases (Li et al., 2007; Boland and Christman, 2008; Sandoval et al., 2019). Hence, the opposing activities of DNMTs, TETs and TDG finely tune methylation dynamics that allow proper cell differentiation and identity acquisition. Besides this function in the CpG methylation cycle, TDG has also been shown to interact physically with the histone acetyltransferase complex CBP and with a number of transcription factors such as p53 and nuclear receptors, including retinoic acid (RA) receptors (RARs), and thereby participates in rendering chromatin accessible to DNA binding factors (Um et al., 1998; Tini et al., 2002; Onabote et al., 2022; Aranda et al., 2023). Hence, TDG is central to mechanisms regulating gene expression during cell-type specification processes, including those triggered by RA.

The RA/RAR pathway exerts pleiotropic effects during development, in particular on cell fate commitment and identity acquisition processes, some of which can be recapitulated *in vitro*. Notably, RA treatment triggers neural differentiation of ESCs as well as of embryonal carcinoma cells (ECCs - Jones-Villeneuve et al., 1982; Okada et al., 2004). Investigating the role of TDG in RA-induced neural differentiation has been previously attempted through *Tdg* inactivation approaches both in mESCs and in mouse primary neural stem cells (NSCs - Cortazar et al., 2011; Wheldon et al., 2014; Steinacher et al. 2019). However, massive cell death occurred in both models early during neural differentiation, precluding an in-depth study of TDG function in neural differentiation processes. To circumvent these limitations, we took advantage of a mouse ECC line called P19 (McBurney, 1993). P19 ECCs derive from epiblast stem cells (EpiSCs) and as such present genic and morphological characteristics of EpiSCs (Han et al., 2013). Furthermore, P19 ECCs can differentiate in the presence of RA (Jones-Villeneuve et al., 1982), and are known as an appropriate alternative model to ESCs and EpiSCs when high cell death is observed (Magnúsdóttir et al., 2013). Using P19 ECCs, we succeeded in establishing a viable *Tdg*-null model cell line that withstands RA treatment, providing us with a unique opportunity to study the role of TDG during cell differentiation. Using this model, we established the impact of *Tdg* inactivation on the transcriptome and demonstrated that TDG is required for balancing cell differentiation towards a neural fate at the expense of a cardiac mesoderm one. During differentiation, *Tdg* inactivation led to changes in nucleosome positioning and to a down-regulation of ATF4-target genes, revealing that TDG is required to maintain ATF4-dependent gene transcription. ATF4, a bZIP transcription factor, confers to cells adaptability to various stress conditions like high levels of ROS, amino-acid (aa) deprivation and endoplasmic reticulum stress, through direct binding to promoters and co-activation by LSD1 and p300/CBP of a large set of genes grouped into the so-called integrated stress response (ISR) pathway (Lassot et al., 2005; Pakos-Zebrucka et al., 2016; Faletti et al., 2021). We finally report that TDG maintains ATF4-target gene expression through regulation of the mTORC1 signaling complex and propose that TDG regulates cell fate acquisition through diverse molecular mechanisms converging on ATF4 activity and which might be relevant to human cancer.

## Materials and Methods

### Cell Culture and treatments

P19 mouse embryonal carcinoma cells (ECCs) were grown and differentiated as described (Sérandour et al. 2012). Briefly, cells were cultured on cell culture plastic dishes in DMEM containing 10% FCS, penicillin and streptomycin. To obtain cell aggregates, 10^6^ cells were grown in bacterial grade plastic dishes with or without 1 μM RA (Sigma-Aldrich, R2625). For mTOR inhibition experiments in undifferentiated ECCs, subconfluent monolayers of cells grown in 6-well plates (10^5^ cells per well) were treated with 20 nM rapamycin (ApexBio, A8167) for 16 hours. To evaluate the impact of mTOR inhibition during RA-induced differentiation, 4×10^4^ cells were seeded in 6-well plates and first treated with 1 μM RA for 24 hours and then with 20 nM rapamycin for an additional 24 hours. Halofuginone (8 nM, Supelco, 32481) or L-proline (0.5 mM, Sigma-Aldrich, P0380) were applied to monolayers of ECCs for 48 hours. The ferroptosis inducer RSL3 (Sigma-Aldrich, SML2234) was applied to monolayers of ECCs for 24 hours. The ferroptosis inhibitor ferrostatin-1 (Fer1, Sigma-Aldrich, SML0583) was applied to cells at 20 μM for 24 hours, in combination with 100 nM RA. MTT assays were run as previously described (Laurent et al., 2022).

### CRISPR-Cas9-mediated *Tdg* knock-out

Knock-out was performed using the *Tdg* mouse Gene Knock-Out kit (Origene, KN317363). This kit contained two gRNAs that targeted the first exon of *Tdg* gene and a donor vector that contained a GFP-Puromycin cassette to facilitate the screening process. Briefly, wt ECCs were seeded in P100 Petri dishes transfected with 1 µg of either gRNA vectors and 1 µg of the donor vector using the JetPEI transfection reagent (Polyplus-transfection). Five passages after the transfection, Puromycin was added to the medium (0.5 µg/ml). After the ninth passage, fluorescence activated cell sorting was carried out to isolate GFP-positive cells (LSR Fortessa X-20, Becton Dickinson). A limiting dilution strategy was then applied to isolate and expand clones from single cells. These clones were analyzed by RT-PCR and PCR on genomic DNA (primers are listed in Supplementary Table 1) to assess for the absence of *Tdg* mRNA and the presence of the GFP-Puromycin cassette respectively.

### CRISPR-Cas9-mediated *Cyp26a1* knock-out

Two guide RNAs (gRNAs) flanking the region of interest were designed using CRISPOR (http://crispor.tefor.net/) and introduced either into pX458 or pX459 (one gRNA, Addgene) at the BbsI (NEB, R0539) site. Wt ECCs were seeded into P100 Petri dishes and transfected for 24 hours with vectors containing the gRNAs, i.e., pX458, carrying a GFP coding sequence, and pX459 with a ratio 1:1.2. GFP-positive cells were isolated using flow cytometry and seeded into a 96-well plate (LSR Fortessa X-20, Becton Dickinson). Clones were analyzed by PCR and knock-out was confirmed by Sanger sequencing.

### MNase assay

For MNase-seq assays, 10-20×10^6^ cells were incubated for 10 min in 1 % formaldehyde in PBS, washed twice and scraped off of plates in ice cold PBS and spun down at 100g for 10min at 4°C. Cells were re-suspended in digestion buffer (50 mM Tris-HCl pH7.6, 2 mM CaCl2, 0.2 % NP40) containing protease inhibitors (complete EDTA-free, Roche) and prewarmed to 37°C for 2 min. Then, 4U of MNase were added to wt ECCs and the reaction mixture was incubated at 37°C for 5 min. For *Tdg*-null ECCs, 2U of MNase were added, due to the limited amount of recovered mono-nucleosomal DNA when using 4U of MNase. The reaction was stopped by adding EDTA to a final concentration of 10 mM. Cells were then incubated for 10 min on ice in SDS lysis buffer (1 % SDS, 50 mM Tris pH8, 10 mM EDTA pH8). An equal volume of TE/RNase buffer (10 mM Tris pH8, 0.1 mM EDTA pH8, 0.4 µg/ml RNase A) was added before incubating at 37°C for 2 hours. A Proteinase K digestion (1 mg/ml) was then performed at 55°C overnight. The mixture was then extracted twice with Phenol/chloroform and the DNA was precipitated 1 hour at −20°C in the presence of glycogen (20 µg). After centrifugation, DNA pellets were washed by ethanol, dissolved in TE buffer (10 mM Tris pH8, 1 mM EDTA pH8) and purified using NucleoSpin Gel and PCR clean-up (Macherey-Nagel). DNA was loaded on a 2 % agarose gel. The mono-nucleosome band was excised and the corresponding DNA was purified using MinElute Gel Extraction kit (Qiagen). Mono-nucleosomal DNA was further analyzed by deep sequencing. MNase-qPCR assays followed the same procedure except that increasing concentrations of MNase were used. After PCR amplification of selected nucleosomes, values were normalized to the amplification values of undigested purified genomic DNA. Primers are listed in Supplementary Table 1.

### Chromatin Immunoprecipitation (ChIP)

ChIP assays of H3K27ac were run as previously described (Sérandour et al., 2012). For TDG and ATF4 ChIP assays, chromatin from wt ECCs was cross-linked using 1% formaldehyde for 10min at room temperature. Cells were rinsed with cold PBS, harvested and lyzed in Farnham lysis buffer (5 mM PIPES pH8, 85 mM KCl, 0.5 % NP40, Protease inhibitor cocktail). Nuclei were pelleted and resuspended in RIPA buffer (1xPBS, 1 % NP40, 0.5 % Sodium deoxycholate, 0.1 % SDS, protease inhibitor cocktail). Chromatin fragmentation was performed by sonicating samples 14 min (30 sec on/off cycles) using a Bioruptor (Diagenode). The fragmented chromatin was incubated at 4°C overnight either with a rabbit polyclonal antibody against mTDG (Gallais et al., 2007), or a rabbit monoclonal antibody against hATF4 (Cell Signaling, D4B8) in RIPA buffer. Complexes were recovered after incubation with 50 μl protein A–conjugated Sepharose bead slurry at 4°C. Beads were then washed four times by LiCl washing buffer (100 mM Tris pH7.5, 1 % Sodium deoxycholate, 1 % NP40, 500 mM LiCl, protease inhibitor cocktail) and twice by TE buffer (10 mM Tris HCl pH8, 1 mM EDTA). Fragments were eluted with extraction buffer (1 % SDS, 100 mM NaHCO3). Cross-linking was reversed by overnight incubation at 65°C and DNA fragments were purified using NucleoSpin Gel and PCR Clean-up columns (Macherey-Nagel).

### RNA preparation, reverse transcription and real-time PCR (qPCR)

RNA was isolated using Trizol reagent (Ambion, Life Technology) according to the manufacturer’s protocol. Reverse transcription was performed using 1 µg of total RNA as template, 200 U of the M-MLV Reverse Transcriptase (Invitrogen) and 250 ng of Pd(N6) random hexamers (Jena Bioscience). Real-time PCR was performed using SYBR Green Master Mix (Bio-Rad) on a Bio-Rad CFX96 machine. All primers were designed with Primer3 (Untergasser et al., 2012) and purchased from Sigma (Supplementary Table 1).

### RNA Sequencing Methodology

Total RNA was extracted using Trizol and RNA Integrity Number (RIN) score were determined with a 2100 Bioanalyzer (Agilent). The protocol for 3’ digital gene expression profiling was carried out as previously described (Chaumette et al., 2022). Library preparation was initiated from 10 ng of total RNA per sample. mRNA poly(A) tails were tagged with universal adapters, well-specific barcodes and unique molecular identifiers (UMIs) during the template-switching reverse transcription process. Barcoded complementary DNAs (cDNAs) from multiple samples were pooled, amplified, and subjected to tagmentation, focusing on enriching the 3’ ends of cDNAs. The resulting library, with fragment sizes ranging from 350 to 800 bp, underwent sequencing in a 100-cycle S2 run on the NovaSeq 6000 system at the Genomics Atlantic platform (Nantes, France). Raw sequencing data were deposited in GEO under accession number GSE262587. On average, each sample yielded approximately 5 million 75 bp single-end reads. Post-sequencing, data demultiplexing was performed using Illumina bcl2fastq. The resulting FASTQ files were analyzed using the 3’ Sequencing RNA Profiling (SRP) pipeline (Charpentier et al., 2021). This methodology aligns with previous DGEseq analyses in various studies (Chaumette et al., 2022; Letellier et al., 2022; Ménoret et al., 2023). The SRP pipeline incorporates cutadapt for read trimming and bwa aln (version 0.7.17) for alignment to the mm10 Mus musculus reference genome and transcriptome. A custom Python script was employed to parse and count UMIs, leading to the generation of a raw matrix containing unique transcript counts. Gene annotation was based on the RefSeq database. The dataset encompassed a total of 26,214 genes, where 3,453 genes had zero reads across all samples, while 22,761 genes had at least one read in at least one sample. The tests for differential expression between two conditions were performed using the R package DESeq2 (Love et al., 2014) version 1.42.0. A gene was declared modulated if it displayed a significant difference between the two indicated conditions with a cut-off fixed at 5.10^−2^ for adjusted P-value (Benjamini-Hochberg). MA-plots show the shrunken log2 fold changes between the indicated conditions, using the adaptive shrinkage estimator from the ashr package (Stephens, 2016). Heatmap in figures 2B and 3F were computed using the ggplot2 R package (Wickham, 2016).

### Library preparation, sequencing, and bioinformatics

All MNase-seq libraries were prepared using the TrueSeq ChIP Sample Prep kit (Illumina) and single-end sequenced using a HiSeq 1500 sequencing system at the “Human and Environmental Genomics” platform (now EcoGeno) (Rennes, France). Raw sequencing data were deposited in GEO under accession number GSE261602. Sequencing reads were mapped to mm8 using Bowtie (Langmead et al., 2009) and processed by SAMtools (Li et al., 2009) to generate bam files followed by wig file generation with MACS (Zhang et al., 2008). Peak calling followed a previously described procedure (Sérandour et al., 2012). Sequencing reads from publicly available datasets used in this study were downloaded from GEO at NCBI and were mapped and treated following the same procedure. For comparison, sequencing data were normalized with respect to the number of reads. To generate heatmaps, except for Fig. 4C which was obtained online with Cistrome (Liu et al., 2011; http://cistrome.org/ap/root), signal matrix was computed using DeepTools (v 3.5.1) computeMatrix tools (Ramírez et al., 2016) and processed using Profileplyr R package (v 1.20.0) (Carroll and Barrows, 2024). To compare the genome-wide distribution of TDG with candidate cis-regulatory elements (cCREs - ENCODE Project Consortium et al., 2020), cCRE coordinates (mm10) from the 2nd version of the registry were downloaded (https://screen.encodeproject.org/index/cversions). CpG island coordinates (mm10) were obtained from UCSC genome browser (https://genome.ucsc.edu/). Transcription start sites (TSSs) from the mouse genome (mm10) were extracted from the Ensembl v102 gtf (https://www.ensembl.org/). Motif enrichment in TDG ChIP-seq peaks was run using the SeqPos tool from Cistrome. The online Dependency Map (DepMap) portal was used in custom analysis mode (https://depmap.org/portal/interactive/custom_analysis) to investigate the correlation between *TDG* gene expression and protein levels in human cancer cell lines, as well as between the occurrence of damaging mutations in the *TDG* gene and growth of human cancer cell lines following CRISPR mediated gene inactivation (Gene effect). Bar graphs (mean +/- SD, n=3 to 6) were generated with GraphPad Prism 5.0 and analyzed by unpaired t-test (*: P<0.05, **: P<0.01, ***: P<0.001, ****: P<0.0001).

### Total cell extract and immunoblot

Total cell extracts were prepared and immunoblotting experiments performed as described previously (Watrin et al, 2014). Rabbit antibodies against U2AF65 (sc-53942) were purchased from Santa-Cruz Biotechnologies, p70S6K (# 9202) and T389p70S6K-Ph (# 9234) from Cell Signalling Technology, and MEIS1 from Abcam (#19867). Rabbit anti-TDG antibodies were already described (Gallais et al., 2007). All antibodies used in this study were affinity-purified.

## Results

### TDG maintains retinoic acid-induced differentiation of ECCs towards neural fate

In order to characterize the dynamics of gene expression during RA-induced ECC differentiation, we first ran a time-course analysis of the transcriptome of P19 ECCs treated with 1 μM RA, by 3’RNA-seq (Supplementary figure 1A). Results indicated that P19 ECCs expectedly lost expression of pluripotency genes upon RA treatment and acquired a posterior fate, as revealed by the rapid extinction of *Otx2* expression and induction of *Cdx1* and *Hoxa1* (Supplementary Fig. 1A). Appearance of posteriorization markers was rapidly followed by the expression of neuromesodermal progenitor (NMP) markers, all of which except *Sox2* dropped at 24 hours of RA exposure while neural progenitor cell (NPC) markers rose markedly (Supplementary Fig. 1A). In parallel, mesoderm progenitor cell (MPC) and paraxial mesoderm markers showed no to low induction (except for *Meox1*) while cardiomyocyte markers were induced (Supplementary Fig. 1A). These data indicate that ECCs mainly engage into a neuroectodermal fate upon RA treatment but a fraction of them adopt a cardiac mesoderm fate. Consistent with this observation, signaling pathways favoring NMP establishment and maintenance as well as mesoderm induction (*i.e.*, Fgf, Wnt and Nodal; Henrique et al., 2015; Edri et al., 2019) were turned down at the time when NPC markers started to be expressed (Supplementary Fig. 1A). Hence, upon RA addition, the P19 ECC system recapitulates key steps of EpiSC differentiation into NPCs and cardiomyocyte precursors and therefore stands as a suitable model for investigating mechanisms involved in cell fate acquisition.

To interrogate the role of TDG in RA-induced differentiation of ECCs, the *Tdg* gene was inactivated by CRISPR/Cas9 in wt P19 ECCs (Supplementary Fig 1B). Transcriptomes were determined by 3’-RNA-seq for both wt and *Tdg*-null ECCs, before and after treatment with RA. Differential gene expression analysis between untreated wt and *Tdg*-null ECCs highlighted 977 differentially expressed genes (DEGs, adjusted *P* value < 0.05, Supplementary Table 2). Gene ontology (GO) enrichment analysis revealed no specific annotation for up-regulated genes and “RNA processing” and “amino acid metabolism” annotations for down-regulated genes (Supplementary Fig. 1C), suggesting that TDG is not involved in the repression of a particular set of genes in the absence of RA. Upon treatment with RA for 48 hours, 700 DEGs were significantly up and 609 down (adjusted *P* value < 0.05) in *Tdg*-null cells compared with wt cells (Fig. 1A, Supplementary Table 3). In particular, *Tdg*-null cells showed a decreased expression of NPC-specific markers like *Pax6* and *Dbx1* and an increased expression of mesoderm markers such as *Acta2*, *Myl9* and *Meox1*, indicating differentiation skewing towards cardiac mesoderm (Fig. 1A). In addition, the RA-inducible *Meis1* gene encoding a transcription factor shared between neural and mesodermal lineages showed similar expression levels in both cell lines 48 hours post RA treatment (Fig. 1A), indicating that *Tdg*-null cells retained the ability to respond to RA. Of note, RA-induced expression of the *Cyp26a1* gene was blunted in the absence of TDG (Fig. 1A and supplementary Fig. 1D). CYP26A1 catabolizes the first step of RA elimination by oxidizing RA into 4-oxoRA and, by doing so, reduces the intracellular levels of RA (Langton and Gudas, 2008). Consistent with a reduced catabolism of RA, RA-induction of *Meis1* expression was triggered by lower concentrations of RA in *Tdg*-null cells than in wt cells (Supplementary Fig. 1E). This higher sensitivity to RA of *Tdg*-null cells was also evidenced phenotypically since RA-induced cell growth as 3D-clusters was triggered by lower RA concentrations in *Tdg*-null cells than in wt ECCs (Supplementary Fig. 1F). In order to investigate whether alterations in RA catabolism could account for the differentiation skewing that we observed in *Tdg*-null cells, we generated *Cyp26a1*-null ECCs (Supplementary Fig. 1G,H). In line with results obtained by treating *Cyp26a1*-null ESCs with high concentration of RA (Russo et al., 2022), RT-qPCR analysis of differentiation markers in RA-treated *Cyp26a1*-null ECCs did not show a reduction in expression of neural markers compared to wt ECCs (Fig. 1B). Thus, reduced CYP26A1 level does not account for the differentiation skewing observed in *Tdg*-null cells. Alternatively, it remains possible that the observed reduced expression of NPC markers in *Tdg*-null cells arises from a mere delay in cell’s response to RA. To test this possibility, we characterized expression levels of these markers 48, 72 and 96 hours after RA treatment by RT-qPCR. Results revealed reduced mRNA levels of the neural markers *Pax6* and *Irx3* and increased mRNA levels of the cardiogenic gene *Wnt11* and the cardiac mesoderm marker *Acta2* in *Tdg*-null cells compared with wt cells at all time points (Fig. 1C). Therefore, the NPC marker reduction and cardiac fate marker increase observed upon *Tdg* inactivation were not due to a delayed gene activation in response to RA but, instead, to a switch from a neural to a cardiac mesoderm fate. In line with this observation and consistent with the role of early NODAL signaling in favoring cardiomyocyte differentiation at the expense of neural differentiation in mESCs (Parisi et al., 2003), expression of *Nodal* as well as of *Lefty1* and *Lefty2*, two genes upregulated by NODAL signaling, showed higher levels in *Tdg*-null than in wt cells shortly after RA addition (Fig. 1D). Finally, GO enrichment analysis of up- and down-DEGs (absolute value of Log2 fold change (FC) > 1 and adjusted P value < 0.05) further established the skewing from neural to mesodermal differentiation in RA-treated *Tdg*-null cells (Fig. 1E). Indeed, down-DEGs were associated with neural developmental processes, whereas up-DEGs were associated with cardiac morphogenetic processes (Fig. 1E). In addition, GO terms that relate to aa transport and metabolism were significantly enriched for down-regulated genes in *Tdg*-null versus wt ECCs treated with RA (Fig. 1E). Collectively, these results indicate that TDG is required for efficient RA-induced neural differentiation and suggest that cell lineage bifurcation and aa-related metabolic processes could be functionally linked in differentiating ECCs.

**Figure 1:**
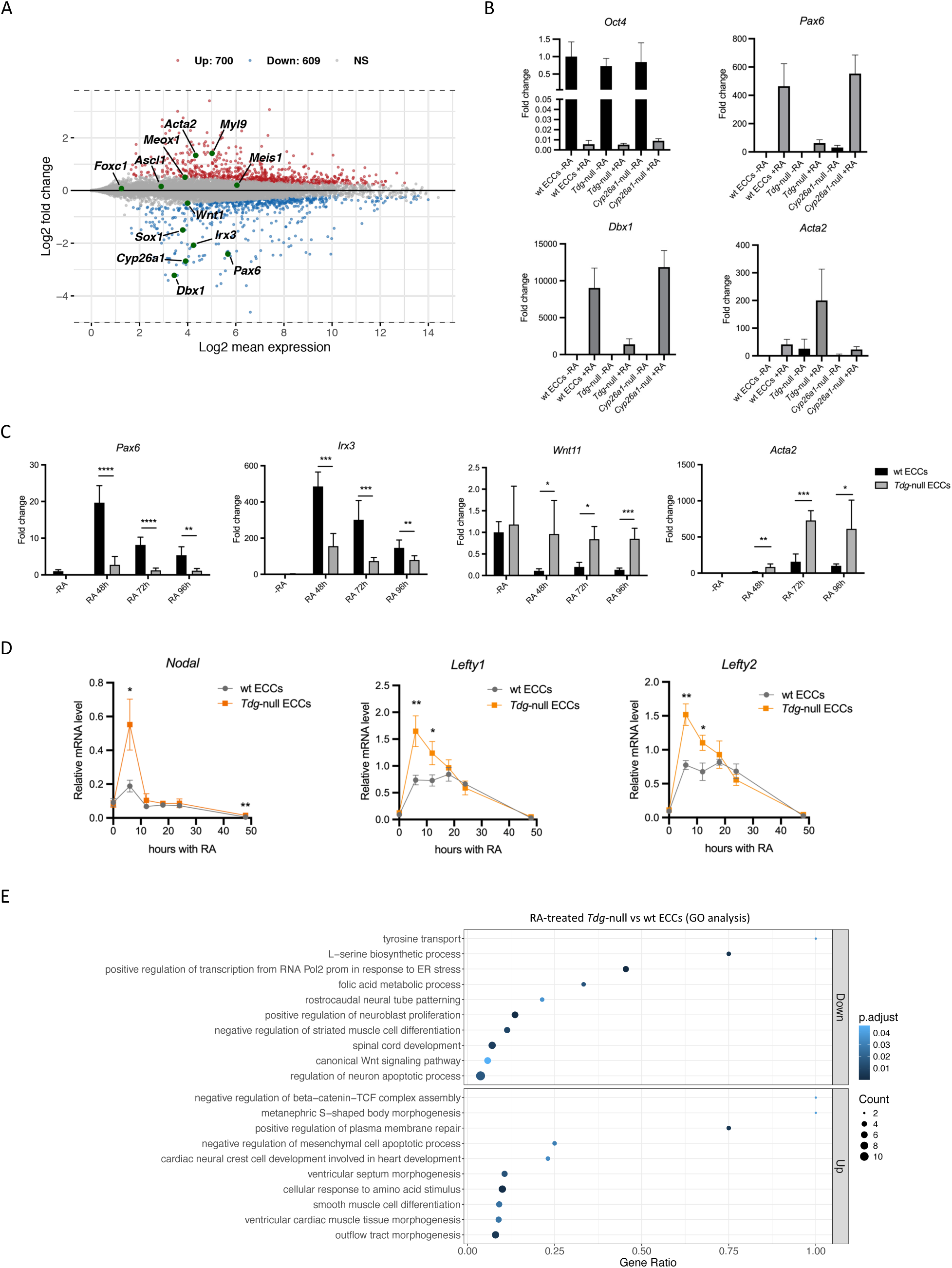
Transcriptomic analysis of wt and TDG-null ECCs undergoing retinoic acid-induced differentiation. (**A**) Log ratio (M) and mean average (A) representation (MA plot) showing differentially expressed genes in 1 μM RA-treated *Tdg*-null ECCs compared to RA-treated wt ECCs grown as monolayers. Up-regulated genes are highlighted in red and down-regulated genes in blue. Specific marker genes are shown in green and their names indicated. (**B**) Quantification by RT-qPCR of mRNA levels of the pluripotency gene *Oct4* and of neural (*Pax6*, *Dbx1*) and cardiac mesoderm (*Acta2*) differentiation marker genes in wt ECCs, *Tdg*-null and *Cyp26a1*-null cells before and after 1 μM RA for 48 hours. For each gene analyzed, expression values are expressed relative to the average expression values observed in untreated wt cells and are shown as “fold change”. (**C**) Quantification of the mRNA levels (RT-qPCR) of the neural markers *Pax6* and *Irx3*, the mesoderm inducer *Wnt11*, and the cardiac mesoderm marker *Acta2* at the indicated time points upon RA treatment of wt and *Tdg*-null ECCS. (**D**) *Nodal*, *Lefty1* and *Lefty2* expression profiles (RT-qPCR) during RA treatment of wt and *Tdg*-null ECCs. (**E**) Gene ontology (GO) enrichment analysis of down- and up-regulated genes described in (A).

### TDG protects ECCs against ferroptosis and regulates ATF4-target genes

*Tdg* inactivation has been associated with high cell death in mouse ESCs and NSCs upon RA treatment (Wheldon et al., 2014; Steinacher et al. 2019). Consistently, we observed that *Tdg* inactivation in ECCs led to increased cell death upon RA treatment, suggesting that TDG also protects ECCs from cell death during differentiation. Indeed, when investigating mitochondrial reductase activity, used as an indirect readout for cell viability through MTT assays, results showed lower viability in monolayers of *Tdg*-null compared to wt ECCs (Supplementary Fig. 2A), suggesting increased cell death in *Tdg*-null cells upon RA treatment. In line with this possibility, a high number of floating dead cells were observed in *Tdg*-null cells treated with 1 μM RA (Supplementary Fig. 2B). Careful examination of transcriptomic data from untreated ECCs did not reveal changes in major components of the apoptotic machinery (data not shown) but instead showed alterations in expression of genes involved in ferroptosis, a non-apoptotic type of cell death that is triggered by an iron-dependent increase in reactive oxygen species (ROS) and in lipid peroxidation that leads to a loss of membrane integrity and eventually to cell death (Jiang et al., 2021). Remarkably, 32% (15 genes, Supplementary Fig. 2C) of the top 47 differentially expressed genes between *Tdg*-null and wt ECCs were previously shown to regulate ferroptosis. These include several genes that protect from ROS accumulation and lipid peroxidation like *Gpx7* (Zhou et al., 2022) and *Mgst1* (Kuang et al., 2021) as well as genes that stimulate ferroptosis like *Upp1* (Lai et al., 2023) and *Phb2* (Yang et al., 2022; Fig. 2A,B; Supplementary Fig. 2C). In addition, the solute carrier genes *Slc3a2* and *Slc7a11*, the products of which form the xc-complex that allows cystine entry into the cell and glutathione synthesis hence protecting cells from ferroptosis (Jiang et al., 2021; He et al., 2023; Fig. 2A), were markedly downregulated in *Tdg*-null versus wt ECCs upon RA treatment (Fig. 2B). Additional time-course study of major antioxidant effectors revealed a RA-dependent increase in *Gpx4*, *Gpx7* and *Mgst1* mRNA levels in wt ECCs that was significantly reduced in *Tdg*-null cells (Fig. 2C). Since expression of a number genes involved in protection against ferroptosis was affected in *Tdg*-null cells, we next evaluated directly the sensitivity of wt and *Tdg*-null ECCs to ferroptosis. In that aim, cells were treated for 24 hours with increasing concentrations of RSL3, a compound inducing ferroptosis by inhibiting the major glutathione peroxidase GPX4 (Yang et al., 2014). Consistent with a dysregulation of genes preventing ferroptosis, *Tdg*-null cells were about 3 times more sensitive than wt cells to RSL3-mediated ferroptosis induction, as determined through mitochondrial activity assays (Fig. 2D). In addition, the ferroptosis inhibitor Fer1 (Dixon et al., 2012) partially restored viability of *Tdg*-null cells treated with 100 nM RA (Supplementary Fig. 2D). These data demonstrate that TDG protects from ferroptosis.

**Figure 2:**
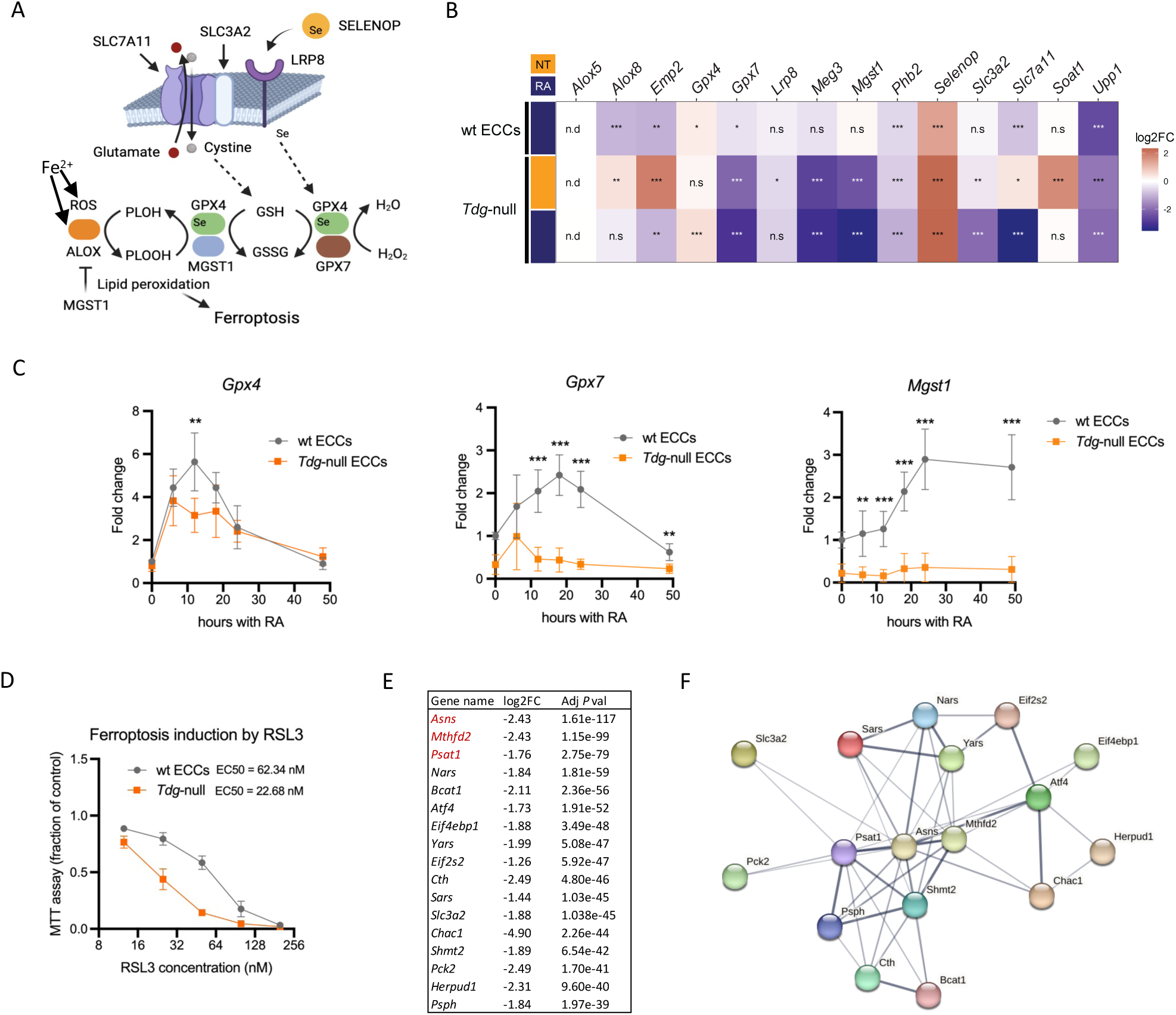
TDG protects cells from ferroptosis. (**A**) Outline of ferroptosis regulation. Ferroptosis is triggered by excessive lipid peroxidation driven by iron-dependent reactions. This can be counteracted by GPX4 and MGST1 in combination with reduced glutathione (GSH). In addition, GPX4 and GPX7, together with GSH, allow to detoxify cells from H2O2, thus lowering ROS production and lipid oxidation. The activity of GPX4 relies on the presence of selenocysteine supplied through degradation of selenoprotein P (SELENOP) upon binding to its receptor LRP8 and endocytosis. Resistance to ferroptosis is also provided by high levels of SLC7A11 and SLC3A2, which allow cystine entry into cells, thus increasing GSH synthesis. (**B**) Log2 fold change (Log2FC) in expression levels of the indicated genes in wt and *Tdg*-null ECCs treated or not (NT) with 1 μM RA for 48 hours. All values are expressed relative to the untreated wt ECCs. (**C**) Quantification of *Gpx4*, *Gpx7* and *Mgst1* mRNA levels (RT-qPCR) during 1 μM RA-response of wt ECCs and *Tdg*-null cells. (**D**) RSL3 dose-response curves in MTT assays. RSL3 EC50 is indicated for both wt and *Tdg*-null cells. (**E**) Log2 fold change (Log2FC) and adjusted *P* value (Adj *P* val) of the 17 most significantly down-regulated genes in 1 μM RA-treated monolayers of *Tdg*-null ECCs compared to RA-treated wt ECCs. (**F**) Network showing functional interactions between the 17 genes and their products shown in (E). The network was drawn using the online tool String (https://string-db.org/). The thickness of the full string network edges indicates confidence in the corresponding supporting data.

It has been established that the ATF4 transcription factor mediates protection against ferroptosis by upregulating the xc-/GPX4 axis (Ahola et al., 2022; He et al., 2023; Swanda et al., 2023). As we observed that the two xc-transporter genes *Slc3a2* and *Slc7a11* were down-regulated in *Tdg*-null cells, we hypothesized that TDG could broadly participate in the transcriptional control of ATF4-target genes. In agreement with this hypothesis, further expression analysis revealed that, strikingly, the 17 most significantly down-regulated genes between RA-treated wt and *Tdg*-null cells all belonged to the so-called integrated stress response pathway (ISR - Pakos-Zebrucka et al., 2023), a cell signaling pathway that depends on ATF4 activity and exerts broad control on protein synthesis in response to metabolic stress (Fig. 2E,F). Remarkably, these 17 downregulated genes are all known direct targets of the transcription factor ATF4 and, consistent with their role in the ISR, are involved in one-carbon metabolism as well as in aa metabolism, including biosynthesis, transport and tRNA charging. Interestingly, expression of the top 3 genes (*i.e.*, *Asns*, *Mthfd2* and *Psat1*, in red Fig. 2E) has been shown to depend on the catalytic activity of TET1 and TET2 in ESCs (Mulholland et al., 2020). As both TDG and TET enzymes act in active DNA demethylation, this observation supports a crucial role of the DNA methylation/demethylation machinery in the control of ATF4 target gene expression in response to RA. Collectively, these data indicate that, in a model recapitulating cell differentiation into NMPs followed by an engagement towards neural or mesoderm fates, TDG prevents ferroptosis and favors the acquisition of a neural fate, possibly by allowing proper expression of ATF4-dependent ISR genes.

### TDG maintains expression of stress-response genes in differentiating ECCs

Since we observed a significant down-regulation of ATF4-target genes in RA-treated *Tdg*-null cells (Fig. 2E), we next addressed the expression dynamics of ATF4-target genes and the role of TDG in their regulation. The time-course response to RA treatment of wt ECC monolayers was analyzed by 3’RNA-seq and revealed that ATF4-target genes exhibited a decrease in mRNA levels during the first 24 hours followed by a recovery between 24 and 48 hours (Fig. 3A). This shared expression pattern was confirmed by RT-qPCR assays for a number of genes (Supplementary Fig. 3A). As an illustration, the solute carrier (*Slc*) family gene members that displayed such expression pattern were ATF4 targets, while ATF4-independent *Slc* genes did not (Supplementary Fig. 3B,C). These changes in gene expression were mirrored by similar variations in acetylation levels of histone H3 lysine 27 (H3K27ac, GEO dataset GSE82314), an active chromatin mark, at ATF4-target gene promoters but not at the ATF4-independent gene *Nampt* (Fig. 3B). Thus, the observed variations in ATF4-target mRNA levels are likely to be regulated transcriptionally, which prompted us to search for genes that are common targets between TDG and ATF4. In order to identify such genes, we reanalyzed publicly available mouse TDG and ATF4 ChIP-seq datasets (GSM1341311 from ESCs and GSM5440974 from cardiomyocytes, respectively). First, we could evidence that promoters from the top down-regulated gene in *Tdg*-null cells (Fig. 2E) are bound by both TDG and ATF4, whereas the *Nampt* promoter was bound by TDG only (Fig. 3C). These data suggest that TDG and ATF4 act directly at promoters in the transcriptional control of ATF4-target genes. Consistently, mRNA levels of ATF4-target genes were decreased in 48h RA-treated *Tdg*-null cells compared to wt ECCs, whereas *Nampt* mRNA levels remained unchanged (Fig. 3D). In addition, acetylation of H3K27 immediately downstream of the transcription start site (TSS) of ATF4-target genes was significantly reduced in RA-treated *Tdg*-null cells compared to wt ECCs, whereas no change was observed for *Atf4* and *Nampt* (Fig. 3E). These data further consolidate the notion that TDG positively regulates ATF4-target genes at the transcriptional level by acting directly at promoters.

**Figure 3:**
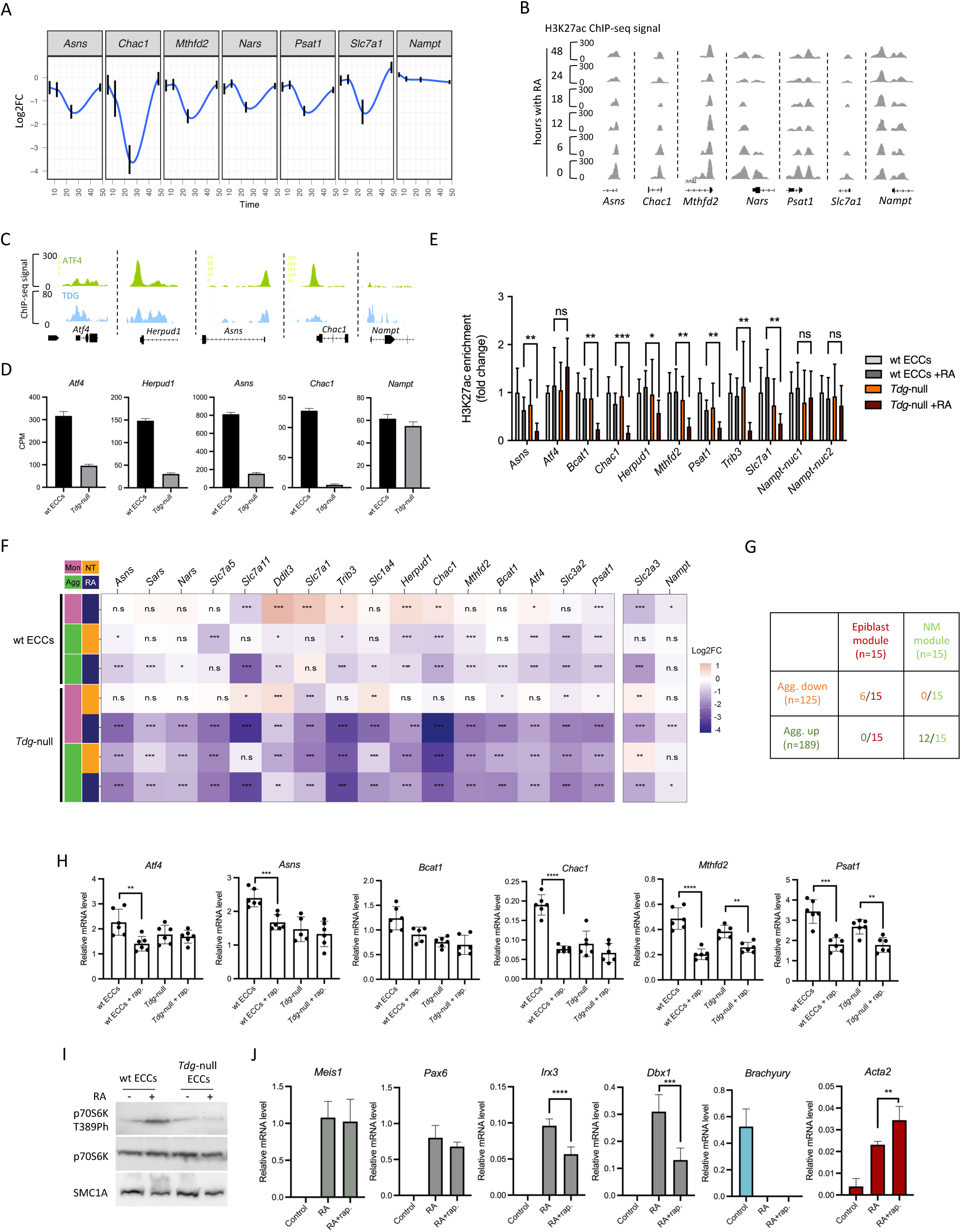
TDG and mTORC1 regulate ATF4-mediated gene transcription during cell differentiation. (**A**) Variations in mRNA levels for the indicated ATF4-target genes and for the unrelated gene *Nampt* (used as a control) extracted from RNA-seq data of a 1 μM RA treatment time-course experiment in wt ECCs. Data are expressed as Log2FC compared to untreated cells. (**B**) Variations in H3K27 acetylation profiles at gene loci shown in (A) during a 1 μM RA treatment time-course experiment in wt ECCs. (**C**) TDG and ATF4 ChIP-seq signals at selected loci. (**D**) Expression levels of the indicated genes in wt and *Tdg*-null ECCs grown as monolayers and treated with 1 μM RA for 48 hours (extracted from RNA-seq data and expressed in counts per million reads - CPM). (**E**) ChIP-qPCR analysis of H3K27ac enrichment immediately downstream of the TSS of ATF4-target genes. For the negative control gene *Nampt*, two different sequences were interrogated (*Nampt*-nuc1 and *Nampt*-nuc2). Values were normalized to the H3K27ac signal in untreated wt ECCs and are expressed as fold change. (**F**) Table displaying Log2 fold change of expression levels for the indicated genes in wt and *Tdg*-null ECCs grown either as monolayers (Mon) or as aggregates (Agg) and treated or not (NT) with 1 μM RA for 48 hours. All conditions are expressed relative to the untreated wt ECCs grown as monolayers. (**G**) Table depicting the overlap between the indicated lists of genes. Agg. up and Agg. down are genes up- or down-regulated by cell aggregation of wt ECCs. Nascent mesoderm (NM) and epiblast gene modules have been previously established (Cheng et al., 2022). (**H**) RT-qPCR analysis of selected ATF4-target gene mRNAs in wt and *Tdg*-null ECCs grown as monolayers and treated or not with 20 nM rapamycin for 16 hours (+ rap.). (**I**) Western blot analysis of the mTORC1 target p70S6K in wt and *Tdg*-null ECC monolayers treated or not with 1 μM RA for 48 hours. Detection of p70S6K phosphorylated on T389 is shown on top, p70S6K in the middle, and the loading control SMC1A on the bottom. (**J**) RT-qPCR analysis of mRNAs encoding the indicated differentiation markers in monolayers of wt ECCs treated or not with 1 μM RA for 48 hours. Twenty-four hours after RA addition, rapamycin was added to the cell culture medium (20 nM; RA+rap.) and cells were further grown for 24 hours.

The observations that the RA-induced formation of 3D clusters of ECCs (Supplementary Fig. 1F) was accompanied by the down-regulation of ATF4-target genes, and that both processes are exacerbated in *Tdg*-null cells, which display increased cell aggregation in response to low concentrations of RA, prompted us to hypothesize that cell aggregation by itself triggers repression of ISR gene expression through the downregulation of ATF4 activity. To test this possibility, wt and *Tdg*-null ECCs were grown as monolayers or as aggregates, treated or not with RA, and the different corresponding transcriptomes were determined by 3’RNA-seq and analyzed. Differential gene expression analysis first revealed that, in accordance with our hypothesis, expression was significantly reduced for 10 out of 16 tested ATF4-target genes when wt ECCs were grown in the absence of RA as aggregates when compared to monolayers (Fig. 3F). When looking for specific cell signatures associated with up- or down-regulated genes in wt ECCs grown as aggregates *versus* monolayers, we evidenced that aggregated cells were primed for pluripotency exit and nascent mesoderm formation. Indeed, 40% (6 out of 15) of the epiblast module genes (Cheng et al., 2022) overlapped with aggregation down-regulated genes, and 80% (12 out of 15) of the nascent mesoderm module genes (Cheng et al., 2022) overlapped with aggregation up-regulated genes (Fig. 3G), indicating that an engagement towards a mesodermal fate associates with a reduction in expression of ATF4-target genes. In the absence of TDG, aggregation had a stronger impact on ATF4-dependent genes since 15 out of these 16 genes showed a significant lower expression together with a higher fold-change amplitude (Fig. 3F). In addition, whereas treatment with RA amplified the effect of cell aggregation on ATF4-target gene expression in wt ECCs, RA did not further reduce expression of most of the tested genes in *Tdg*-null cells (Fig. 3F). Importantly, aggregation had no impact on the expression of the two non-ATF4-target genes *Slc2a3* and *Nampt* (Fig. 3F). Collectively, these data indicate that TDG positively regulates ATF4-target genes in aggregates and that the transient RA-mediated down-regulation of these genes observed in RA-treated monolayers is probably initiated through the 3D reorganization of cells in culture (Supplementary Fig. 3D). In line with these results, we propose that 3D growth of ECCs *per se* induces a transient metabolic stress state that triggers a TDG-dependent ATF4-mediated adaptative response.

ATF4 activity can be tuned by distinct molecular pathways that lead to expression of overlapping sets of ATF4-dependent ISR genes. Cell adaptation to metabolic stress through the ISR relies in part on the stimulation of the *Atf4* mRNA translation by the GCN2 pathway (Harding et al., 2000). In addition, upon growth factor-induced proliferation, *Atf4* mRNA stability and translation are sustained by the mechanistic target of rapamycin complex 1 (mTORC1) in mouse embryonic fibroblasts and various cancer cell lines (Park et al., 2017; Torrence et al., 2021). Notably, ATF4 activation by mTORC1 upregulates a subset of ISR genes that are involved in aa uptake and synthesis as well as in tRNA charging (Torrence et al., 2021). To determine whether one or both of these pathways control ATF4-target gene expression in ECCs, we selectively modulated their activity by a pharmacological approach. First, the GCN2 pathway was investigated either using Halofuginone, a compound that blocks the prolyl-tRNA synthetase, which mimics aa starvation and hence activates GCN2, or using an excess of L-Proline, which depletes cells of uncharged tRNAs thus lowering GCN2 activity (D’Aniello et al., 2015). As shown in Supplementary Fig. 3E, and contrary to what has been observed in ESCs (D’Aniello et al., 2015), Halofuginone did not increase *Asns* or *Psat1* mRNA levels whether in wt or in *Tdg*-null ECCs. In the case of an active GCN2 pathway, L-Proline would induce a decrease in *Asns* and *Psat1* mRNA levels in cells. However, the observed decrease was not significant (Supplementary Fig. 3E). These results indicate that the GCN2 pathway plays no role in the expression of these two ATF4-target genes in ECCs, under these conditions. Next, we addressed the involvement of the mTOR pathway by treating wt and *Tdg*-null ECCs with the mTORC1 inhibitor rapamycin and measuring expression of ATF4-target genes by RT-qPCR assays. As shown in Fig. 3H, all but one ATF4-target genes exhibited reduced expression upon rapamycin treatment in wt cells, indicating that mTOR activity is indeed required for proper expression of ATF4-target genes. Further, these genes showed no to small reduction in expression upon rapamycin treatment in *Tdg*-null cells (Fig. 3H), suggesting that TDG is necessary to maintain sufficient levels of mTORC1 activity. The requirement of TDG for mTORC1 activity was confirmed by western blot analysis that revealed reduced levels of the phosphorylated form of p70S6K, a direct target of mTORC1 and a marker of its activity, in *Tdg*-null cells *versus* wt ECCs (Fig. 3I). In order to investigate how TDG regulates mTORC1 activity, transcriptomic data were interrogated and revealed that known activators of mTORC1 (*i.e*., *Mgst1*, *Gpx7*, *Rpl22l1* and *Slc7a5*) showed reduced expression levels in *Tdg*-null compared to wt ECCs (Supplementary Fig. 3F). In addition, the mTORC1 inhibitor encoding gene *Sesn1* was down-regulated in RA-treated wt ECCs but not in *Tdg*-null ECCs (Supplementary Fig. 3F). Knowing that sestrins (SESN1 and SESN2) are leucine sensors that lose their ability to inhibit mTORC1 when bound to leucine (Wolfson et al., 2016; Cangelosi et al., 2022), the expected decrease in leucine uptake in *Tdg*-null cells through reduced SLC7A5 levels could explain the lower mTORC1 activity detected in *Tdg*-null cells. In addition, reduced mTORC1 activity upon decreased cystine influx through SLC7A11 has been shown to sensitize cells to ferroptosis (Zhang et al., 2021). Hence, TDG appears to be a key regulator of mTORC1 activity in differentiating cells.

The activation of the mTOR pathway observed upon RA treatment (Fig. 3I) and the impact of its inhibition on ATF4-target gene expression (Fig. 3H) strongly suggest that mTOR pathway activity participates in cell differentiation induced by RA. To directly evaluate this possibility, wt ECCs were treated with rapamycin 24 hours after differentiation had been initiated by RA, and expression levels of differentiation markers were determined by RT-qPCR. Whereas expression of the mixed-lineage marker *Meis1* and the early neural marker *Pax6* were not affected by rapamycin treatment, late neural markers like *Irx3* and *Dbx1* were down-regulated, and the cardiac mesoderm marker *Acta2* increased (Fig. 3J).

Collectively, these data reveal that TDG ensures a sufficient level of mTORC1 activity to sustain the transcriptional activity of ATF4-regulated genes and that the mTOR pathway is essential for proper cell fate choice in RA-treated ECCs.

### TDG is engaged at ATF4-target gene promoters and regulates nucleosome positioning

To gain further insight into the role of TDG in the control of gene expression at genome scale, we next compared the genome-wide distribution of TDG in ESCs with candidate cis-regulatory elements (cCREs) of the mouse genome defined by DNase1 accessibility and enrichment in H3K27ac and H3K4me3 (ENCODE Project Consortium et al., 2020). Among the 49,699 identified TDG peaks, 23.5 % (n=11,666) were located at promoters, 20.7 % (n=10,275) mapped within proximal enhancers and 23.5 % (n=11,663) to distal enhancers (Fig. 4A). In agreement with a functional relationship between TDG and ATF4 at promoters, most TDG-bound promoters were also ATF4-bound (Fig. 4A). In addition, 3.4 % (n=1,675) overlapped with strong CTCF binding sites in both ESCs (GSE98671) and ECCs (GSE103198-Fig. 4A). Consistent with a high prevalence of CpG islands (CGIs) at promoters, 69 % (8,049 out of 11,666) of TDG sites overlapping with promoters were included in CGIs (Fig. 4B). Further examination of TDG engagement at CGIs showed that TDG binds to almost all CGIs (Fig. 4C), suggesting that TDG engagement in chromatin depends on CpG density. Search for transcription factor DNA binding motifs within the TDG peaks identified the Sp1 motif (5’-CCCCGCCCC-3’), consistent with an engagement of TDG at CpG-rich sequences, and the CTCF motif as highly enriched (Fig. 4D). In order to unveil any correlation between transcription factor binding at promoters and gene expression levels, genes were next ranked according to their expression in untreated wt ECCs, and the binding of TDG, ATF4 and CTCF around TSSs was analyzed. As shown in Fig. 4E, TSSs of highly expressed genes were found to be associated with both TDG and ATF4 binding and the presence of CGIs, whereas TSSs of low expressed genes were not (Fig. 4E). On the contrary, CTCF showed low signal at TSSs (Fig. 4E). Further, CTCF was not found at TSSs of ATF4-target genes down-regulated by TDG depletion, except for *Mthfd2* in ESCs (Supplementary Fig. 4A). This strongly suggests that CTCF is not directly involved in TDG-mediated regulation at TSSs of ATF4-target genes, both in ECCs and ESCs since CTCF showed a similar distribution in these two cell types (Supplementary Fig. 4B). Consistent with data shown in Fig. 3, down-regulated genes in RA-treated *Tdg*-null cells could be distinguished from up-regulated genes by their association with TDG and ATF4 binding within +/- 100 bp of their TSS (p=0.0014, exact Fisher test) whereas such a difference was not observed for RA-treated wt ECCs (p=0.3645, exact Fisher test). These data suggest that TDG positively regulates ATF4-target genes through direct binding at or near their TSSs.

**Figure 4:**
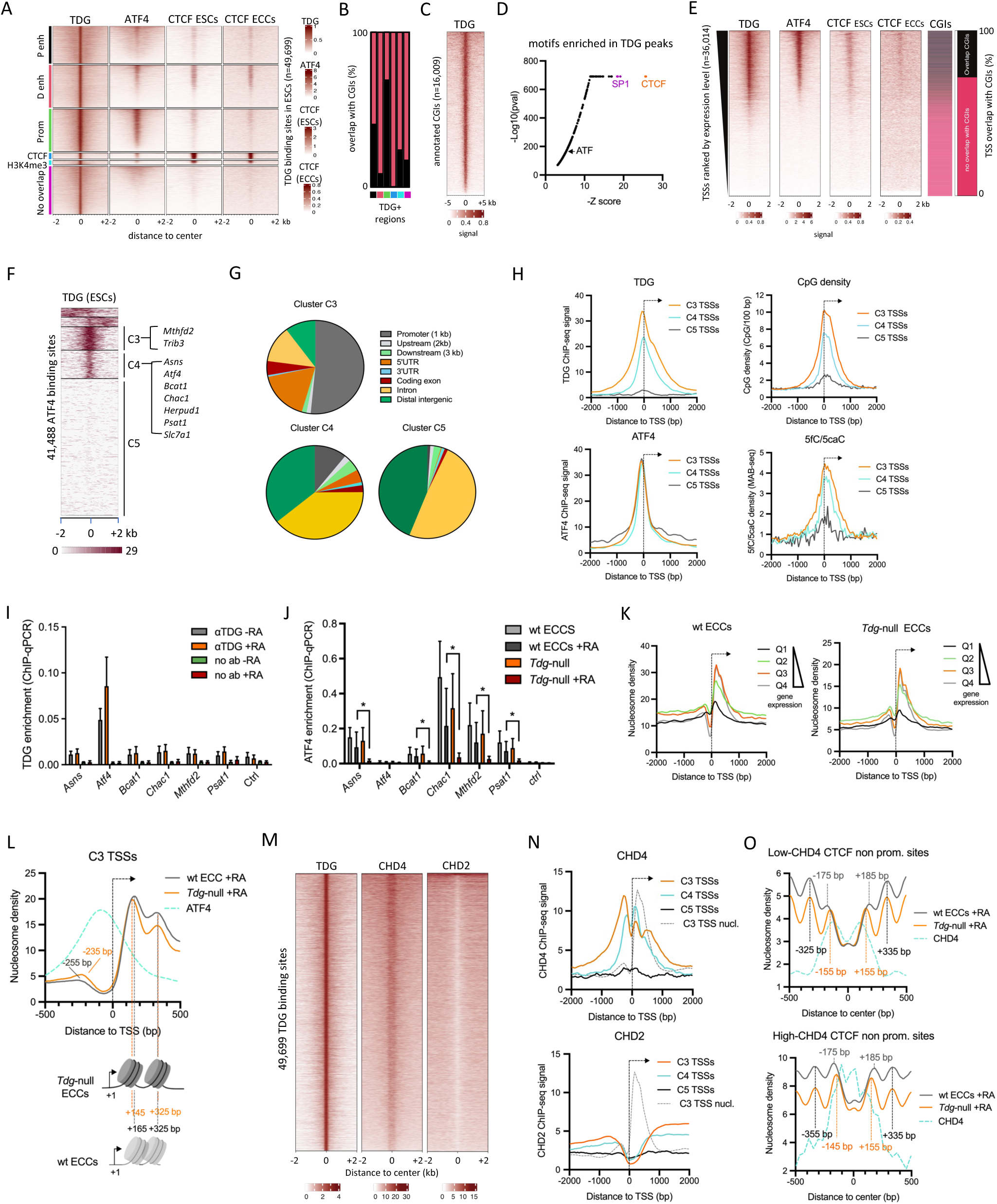
TDG binds active gene promoters and regulates nucleosome positioning. (**A**) Heatmaps of TDG, ATF4, and CTCF ChIP-seq signals in clusters of TDG binding sites overlapping or not with cCREs. The signals are centered on the middle of the TDG peaks (P enh: proximal enhancers, D enh: distal enhancers, Prom: promoters, CTCF: CTCF binding sites, H3K4me3: cCREs marked by H3K4me3 only, No overlap: TDG binding sites not overlapping with cCREs). (**B**) Fraction of TDG binding sites overlapping with CGIs and categorized by their overlap with different types of cCREs. Same color code as in (A). (**C**). Heatmap of TDG ChIP-seq signal in mESCs at annotated CGIs. (**D**) Enrichment in transcription factor motifs at TDG binding sites. (**E**) Heatmap of TDG, ATF4 and CTCF ChIP-seq signals centered on the TSSs of genes ranked by their expression level in wt ECCs. TSSs embedded in CGIs are indicated by a dark gray bar and others by a pink bar on the right side of the heatmaps. (**F**) Clustering heatmap of ATF4 binding sites according to their enrichment in TDG in mESCs. (**G**) Genomic distribution of ATF4 binding sites from clusters C3, C4 and C5 as defined in (F). (**H**) Average profiles of CpG density, 5fC/5caC-seq (MAB-seq) signal, and TDG and ATF4 ChIP-seq signals centered on the oriented TSS (arrow) of genes associated with ATF4 binding (ATF4 ChIP-seq peak within -/+ 500 bp of a single TSS) and included in clusters C3, C4 and C5. (**I**) ChIP-qPCR assay of TDG at ATF4-target gene promoters in wt ECCs. (**J**) ATF4 ChIP-qPCR assay at its target gene promoters in wt ECCs. (**K**) Average profiles of MNase-seq signal at the TSS of genes ranked by quartiles of expression in untreated wt and *Tdg*-null ECCs. (**L**) Average profiles of MNase-seq signal around C3 TSSs in wt and *Tdg*-null ECCs treated with 1 μM RA. The ATF4 ChIP-seq profile is also shown (dashed blue line). Average positions of nucleosome dyads relative to the TSS are indicated. (**M**) Heatmap representation of TDG, CHD4, and CHD2 ChIP-seq signals at TDG binding sites in mouse ESCs. (**N**). Average CHD4 (top panel) and CHD2 (bottom panel) ChIP-seq profiles around C3, C4 and C5 TSSs. The nucleosome density profile around C3 TSSs is also shown. (**O**) Average profiles of MNase-seq signal at non-promoter CTCF binding sites common between ESCs and ECCs in RA-treated wt and *Tdg*-null ECCs and grouped as CHD4-low (top panel) or CHD4-high (bottom panel). The CHD4 ChIP-seq profile has been added in each panel (dashed blue line). The values of CHD4 ChIP-seq signal at CHD4-high CTCF sites have been reduced to fit within the range of nucleosome density values. Average positions of nucleosome dyads relative to the CTCF binding sites are indicated.

To further characterize the functional relationship between ATF4 and TDG engagement genome-wide, we clustered all ATF4 binding sites according to the TDG ChIP-seq signal intensity from ESCs (Fig. 4F). Due to the off-centered distribution of TDG in clusters 1 and 2 (Fig. 4F), these two clusters were not further analyzed. Whereas ATF4 binding sites that were not bound by TDG were mostly intronic and intergenic (cluster C5), sites that were highly enriched in TDG (cluster C3) were mostly associated with promoters and 5’UTRs, reminiscent of CGI distribution along the genome (Fig. 4G). Accordingly, C3 TSSs (n=3,010) showed high TDG and ATF4 binding and high CpG density, whereas C5 TSSs (n=347) showed high ATF4 binding but low CpG density and low TDG engagement (Fig. 4H). Cluster C4 TSSs (n=940) showed high ATF4 binding but intermediate engagement of TDG and lower CpG density than C3 TSSs (Fig. 4H). Similar to TDG distribution patterns, 5fC/5caC accumulation which reflects active turnover of 5mC was higher in C3 than in C4 and almost absent from C5 regions (Fig. 4H), as determined by methylation assisted bisulfite sequencing (MAB-seq) in *Tdg* knock-down ESCs (GEO dataset GSE62631; Neri et al., 2015). Importantly, TDG engagement at *Atf4* and selected ATF4-target promoters was independently confirmed by ChIP-qPCR experiments performed in wt P19 ECCs (Fig. 4I), indicating that TDG regulates ATF4-target gene expression likely through direct binding to their promoter in wt ECCs. Next, we interrogated ATF4 binding to the same selected ATF4-target gene promoters by ChIP-qPCR. Except for *Atf4* itself, data evidenced a marked engagement of ATF4 at all tested promoters as well as a significant decrease in ATF4 binding in RA-treated *Tdg*-null ECCs, consistent with a lower activity of the corresponding genes (Fig. 4J). Together, these results indicate that TDG engagement and activity at ATF4-bound TSSs facilitate gene transcription.

In order to interrogate a putative role of TDG on chromatin organization at ATF4-bound promoters, genome-wide nucleosome maps were generated through MNase-seq experiments that were performed on wt and T*dg*-null ECCs treated or not with RA for 48 hours. First, TSSs were ranked in quartiles of expression in RA-untreated wt and *Tdg*-null ECCs. As previously observed in mESCs (Voong et al., 2016), average profiles of nucleosome occupancy showed a positive correlation with gene expression in both cell lines (Fig. 4K). Nucleosome occupancy around TSSs of genes from cluster C3, cluster C4 and cluster C5 followed this correlation with high nucleosome occupancy around C3 and C4 TSSs being associated with high gene expression and low nucleosome occupancy around C5 TSSs being associated with low gene expression (Supplementary Fig. 4C,D). In addition, a slight decrease in nucleosome occupancy at C3 and C4 TSSs upon RA treatment was observed in both cell lines, an effect that was more pronounced for C3 TSSs (Supplementary Fig. 4C). This observed reduction in nucleosome occupancy was further confirmed by MNase-qPCR targeting the +1 nucleosome of selected ATF4-target genes (Supplementary Fig. 4E). Interestingly, and although nucleosome density did not seem to vary between wt and *Tdg*-null ECCs, nucleosome positioning was markedly altered around the TSSs of C3 genes where the dyad of −1 and +1 nucleosomes appeared shifted towards the TSS by 20 bp in RA-treated *Tdg*-null cells compared to wt ECCs (Fig. 4L). Such a shift was also detected for C4 TSS +1 nucleosomes, whereas C5 genes did not show well-defined nucleosome positions around TSSs (Supplementary Fig. 4F). Examination of ATF4 distribution around C3 and C4 TSSs further indicated that ATF4 binds to the nucleosome depleted region (NDR) of its target genes (Fig. 4L, Supplementary Fig. 4F). Overall, these data strongly suggest that TDG regulates the position of nucleosomes flanking the NDR at promoters of active ATF4-target genes, a process that might favor ATF4 engagement. Given the impact of the +1 nucleosome position and of the organization of a promoter NDR on the transcriptional activity of genes (Bai and Morozov, 2010; Abril-Garrido et al., 2023), these observations support a direct role of TDG in the regulation of ATF4-target gene activity. The observed shift in nucleosome positioning in the absence of TDG is likely to reflect a change in the activity of a remodeler such as CHD4, which is known to associate with nucleosome positioning at active promoters and CTCF binding sites in mouse ESCs (de Dieuleveult et al., 2016; Clarkson et al., 2019). To investigate a putative functional relationship between TDG and CHD4, we next compared the distribution of TDG, CHD4 (GSM1581292, mouse ESCs) and the related remodeler CHD2 (GSM1581290, mouse ESCs), which has been shown to be enriched in gene bodies in correlation with the transcription elongation mark H3K36me3 (de Dieuleveult et al., 2016), around TDG-bound sites in mouse ESCs. Results showed that CHD4 was bound to a large fraction of TDG sites whereas CHD2 showed no binding at these sites (Fig. 4M). Enrichment profiles of CHD4 around C3, C4 and C5 TSSs were consistent with a role of CHD4 in nucleosomal organization around C3 and C4 TSSs since the level of CHD4 peaks at nucleosome positions, as shown for C3 TSSs (Fig. 4N). On the contrary, CHD2 was not enriched at these nucleosome position (Fig. 4N).

Since CTCF-bound sites are known to have well positioned nucleosomes (Clarkson et al., 2019), we next interrogated the role of TDG in nucleosome positioning at non-promoter CTCF binding sites that are common between ESCs and ECCs (n=29,207). Similar to the situation at TSSs, the positioning of nucleosomes directly flanking CTCF binding sites was altered in *Tdg*-null cells compared to wild-type cells in both RA-treated and control conditions, whereas the position of more distant nucleosomes remained unaltered (Supplementary Fig. 4G). Whereas the dyad of these CTCF-flanking nucleosomes was positioned 175 to 185 bp away from the CTCF binding sites in wt ECCs, as shown for CTCF sites in ESCs (Clarkson et al., 2019), its distance to CTCF sites was lowered by 20 to 30 bp in *Tdg*-null ECCs compared to wt ECCs, at either low or high CHD4 sites (Fig. 4O). Consistent with a role of CHD4 in mediating TDG role in positioning these CTCF-flanking nucleosomes, CHD4 was found to preferentially bind next to these two nucleosomes (Fig. 4O). Collectively, these data indicate that TDG is involved in the precise positioning of nucleosomes flanking the TSS of ATF4-target genes and CTCF-bound sites by regulating the recruitment and/or activity of chromatin remodelers.

### *TDG* expression correlates with expression of ATF4-target genes in human

Our data show that TDG plays a role in the regulation of the ATF4/ISR pathway in mouse ECCs. To investigate whether such regulatory function is conserved in humans and relevant to pathologies, we interrogated a putative correlation between the expression levels of *TDG* and *ASNS* (as a paradigmatic stress response gene) expression, using gene expression data from 37 cohorts of patients that cover a wide range of cancer types and are publicly available through the UCSC Xena browser (https://xenabrowser.net/). This analysis revealed that a handful of cancer types exhibit a strong positive association (Pearson’s rho ≥ 0.4, p ≤ 0.05) between *TDG* and *ASNS* expression levels (Fig 5A). The highest correlation was found for patients affected by pediatric brain tumors (Children’s Brain Tumor Tissue Consortium-CBTTC, 854 tumors from 33 different types) where *TDG* expression was of adverse prognosis (p = 1.12E-6), as was that of *ASNS* (p = 2.48E-13). Further examination of *ASNS* and *TDG* expression in the most represented tumor types within pediatric brain tumors showed variable expression and correlation levels of these two genes, with the highest correlation being observed for Low grade glioma (Fig. 5B,C). Interestingly, activation of the *ASNS* gene by ATF4 in asparagine-depleted acute lymphoblastic leukemia cells was shown to be counteracted by DNA methylation (Jiang et al., 2019), suggesting that expression of this ISR gene is repressed by DNA methylation in human and is activated by oxidation and/or erasure of the methylation mark. Thus, we next tested the existence of such an inverse correlation between DNA methylation and *ASNS* expression in the Cancer Genome Atlas (TCGA) lung cancer cohort, in which correlation between *TDG* and *ASNS* expression was also high (rho = 0.543, p = 1.285E-87, Fig. 5A). While most CpGs located immediately upstream of the *ASNS* gene were unmethylated in patient cells (not shown), cg25906151 methylation did exhibit an inverse correlation with both *ASNS* and *TDG* expression (Fig. 5D,E), suggesting that increased DNA methylation at this CpG negatively impacts on *ASNS* expression in lung cancer patients. Consistent with the positive role of TDG in the regulation of ATF4-target genes in mouse ECCs, the expression of other ATF4 target genes such as *MTHFD2* and the aa transporters *SLC7A1* and *SLC7A5* also showed a positive correlation with *TDG* expression in lung cancer patients (Fig. 5F). Conversely, expression of the glucose transporter gene *SLC2A3*, which is not an ATF4 target in mouse, did not show any correlation with that of *TDG* (Fig. 5F). Finally, *TDG* expression was negatively correlated with that of the mTOR pathway inhibitor *SESN1*, whereas a positive correlation was observed with *RPL22L1*, a positive regulator of mTOR signaling (Fig. 5F, Supplementary Fig. 3F). In the TCGA lung cancer cohort, expression correlation with the selected genes was higher for *TDG* than for *ATF4* expression (Fig. 5G). These analyses confirmed that TDG levels impact on ATF4 and mTOR pathways at gene expression level and revealed that the TDG-ISR axis is likely involved in the pathogenesis of some human cancer types In line with a putative implication of TDG in the control of ferroptosis sensitivity in human cancer cells, analysis of the correlation between *TDG* expression and protein levels through interrogating proteomics data obtained in cancer cell lines (https://depmap.org/portal/interactive/custom_analysis) revealed that highly significant correlations were observed for ferroptosis-related proteins (Fig. 5H), such as the pro-ferroptotic proteins MVP, FOSL1 and CTSB (Kuang et al., 2020; Shao et al., 2023; Xia et al., 2024), or the anti-ferroptotic protein CD44 (Bian et al., 2023). The role of TMPO in ferroptosis has not been reported yet but the oncogenic TMPO antisense RNA 1 (TMPO-AS1) is part of a long non-coding RNA signature of ferroptosis and positively regulates TMPO production (Li et al., 2020; Yao et al., 2021). Such positive (TMPO) or negative (MVP) correlation with TDG was also observed at the mRNA level in tumors from the TCGA lung cancer cohort (Supplementary Fig. 5A). When interrogating gene dependencies (DepMap CRISPR screen project Chronos) in cancer cell lines harboring *TDG* damaging mutations compared to other cell lines, we found that only two genes showed a significant correlation with *TDG* mutations: the ferroptosis resistance gene *FBXL5* (Liu et al., 2024) and the mTOR inhibitor gene *GNAO1* (Kim et al., 2024 - Fig. 5I, Supplementary Fig. 5B). Consistent with our results in mouse ECCs showing that *Tdg*-null cells are primed for ferroptosis, the growth of cells with damaging mutations of *TDG* was more sensitive than other cells to *FBXL5* disruption (Fig. 5I, Supplementary Fig. 5B). Also, as *Tdg*-null ECCs showed lower mTORC1 activity, the growth of *TDG* mutant cancer cells was positively impacted by *GNAO1* disruption (Fig. 5I, Supplementary Fig. 5B). In lung cancer patients, the higher expression of *FBXL5* in tumor cells expressing low levels of *TDG* and low levels of *SLC7A11* (Fig. 5J) is anticipated to protect cells from ferroptosis, hence allowing for tumor growth. Extending our findings to humans, these data indicate that TDG is likely to be involved in ATF4-mediated gene regulation, as well as in the control of ferroptosis and of the mTOR pathway in cancer patients. Beside its essential role in cell homeostasis, the ATF4-controlled ISR also stands crucial for the growth of certain cancer types (Tian et al., 2021; Lines et al., 2023). Hence, our data support a conserved role of TDG in the control of ISR genes and suggests and involvement of dynamic DNA methylation processes in cancer cell growth through the regulation of ATF4-dependent genes.

**Figure 5:**
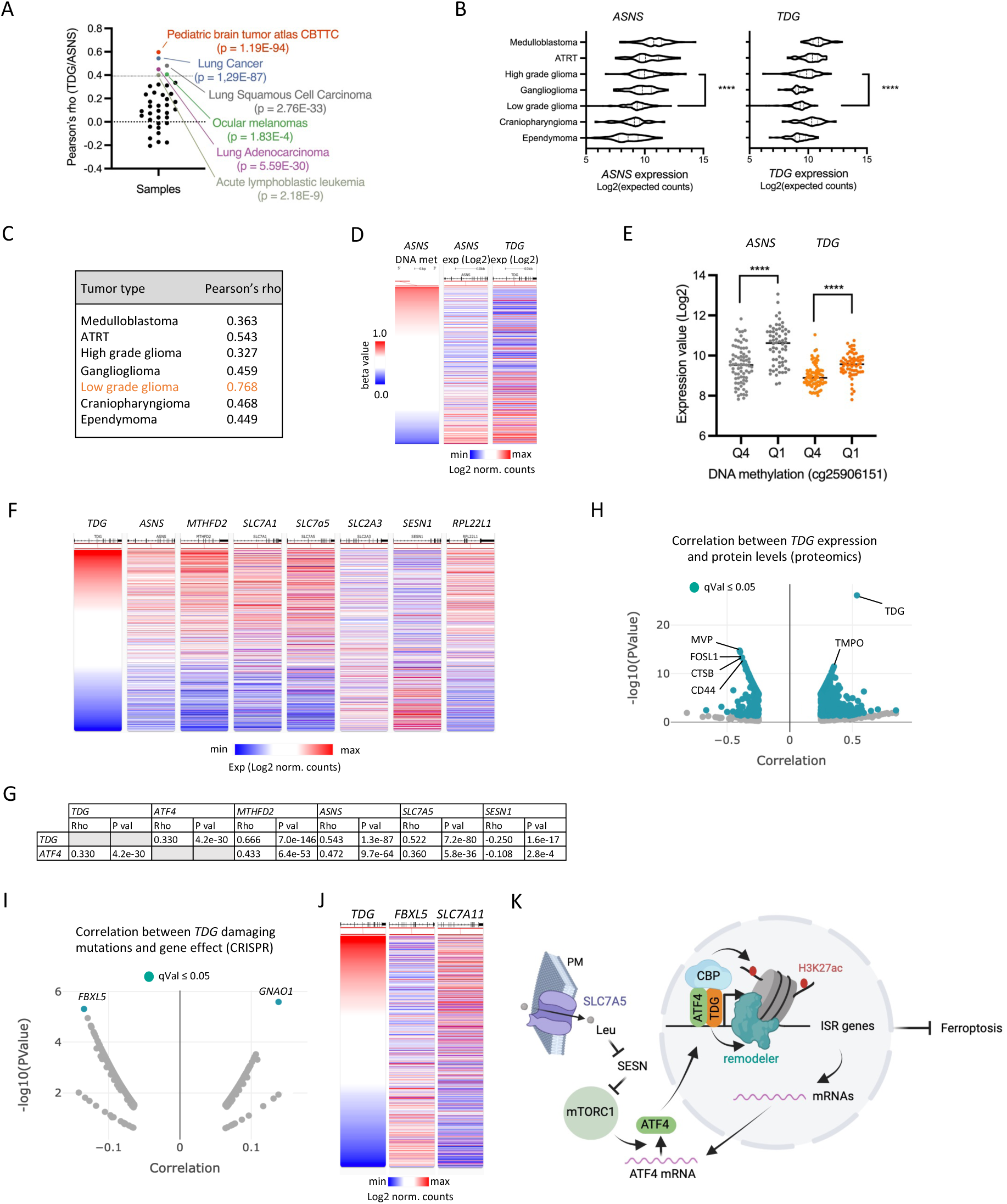
*TDG* and ATF4-target gene expression correlate in human cancer. (**A**) Pearson’s correlation between *TDG* and *ASNS* expression levels in patient tumors from 37 TCGA cohorts. (**B**) Violin plots of *ASNS* and *TDG* expression levels in pediatric brain tumors (ATRT: Atypical Teratoid Rhabdoid Tumor). (**C**) Pearson’s correlation between *TDG* and *ASNS* expression levels in pediatric brain tumors. (**D**) Heatmap representation of cg25906151 methylation levels and *ASNS* and *TDG* expression levels in lung cancer samples (n=812; high methylation/expression: red, low methylation/expression: blue). (**E**) Expression levels of *TDG* and *ASNS* in lung cancer samples classified by quartiles of cg25906151 methylation (Q1: lowest DNA methylation, Q4: highest DNA methylation). (**F**) Heatmaps showing the expression levels of the indicated ISR genes and mTOR pathway genes in lung tumor samples (TCGA lung cancer cohort, n=1,129) ranked according to *TDG* expression levels. (**G**) Pearson’s correlation between expression levels of either *TDG* or *ATF4* and selected genes in lung cancer patients. Rho and P values are indicated for each combination. (**H**) Correlation between *TDG* expression (mRNA levels) and protein levels in human cancer cell lines (DepMap custom analysis, proteomics). Proteins with a correlation qVal ≤ 0.05 have been highlighted in green. (**I**) Correlation between the occurrence of *TDG* damaging mutations and the gene effect observed in CRISPR experiments (DepMap Public 24Q2+Score, Chronos). Gene effects with a correlation qVal ≤ 0.05 have been highlighted in green. (**J**) Heatmaps showing the expression levels of the indicated genes in lung tumor samples (TCGA lung cancer cohort, n=1,129) ranked according to *TDG* expression levels. (**K**) Working model depicting ISR gene regulation by TDG. ATF4 and TDG co-binding at ATF4 target gene promoters is associated with H3K27 acetylation by CBP and +1 nucleosome positioning by chromatin remodeler(s), leading to expression of ISR genes, which include aa transporter genes such as *Slc7a5*, and favoring resistance to ferroptosis (PM: plasma membrane). The resulting leucine (Leu) influx stimulates mTORC1 activity by inhibiting sestrins (SESN). This, in turn, favors *Atf4* mRNA stability and translation, sustaining ISR gene activity.

## Discussion

Deciphering the role of TDG in cell lineage commitment has been hampered so far by the cell lethality that is caused by RA treatment in *Tdg* knock-out mESCs as well as in *Tdg* knocked-down primary neural stem cells (Wheldon et al., 2014; Steinacher et al. 2019). Here, we describe the establishment of a *Tdg* knock-out murine cell model derived from wt ECCs. These cells proliferate *in vitro* and maintain viability upon RA-induced differentiation, therefore representing a relevant substitute model for EpiSCs that allowed us to investigate the molecular function of TDG in gene control and cell fate commitment.

In the course of this study, we could evidence that TDG is required for efficient neural differentiation, as *Tdg*-null cells underwent differentiation skewing towards a cardiac mesoderm fate when treated with RA. Similarly, a transient *Tdg* knock-down followed by a recovery period before embryoid body differentiation of mESCs led to an increase in cardiac mesoderm formation, possibly through a down-regulation of repressors of mesoderm differentiation (Aranda et al., 2023). Differentiation skewing towards cardiac mesoderm has also been observed *in vivo* in TET mutants (Li et al., 2016), suggesting a common requirement of various components of the active DNA demethylation machinery for neural differentiation. Consistent with the idea that DNA demethylation associates with neural differentiation, the knock-out of the DNA methyltransferase gene *Dnmt3b* in mESCs inversely skews differentiation towards neurectoderm (Lauria et al., 2023). Altogether, these observations converge to a model where mechanisms controlling DNA methylation dynamics are important components of cell fate commitment.

Although a number of differentially methylated regions (DMRs) observed in EpiSC-like cells derived from *Dnmt3b*-KO ESCs (Lauria et al., 2023) were found to overlap with the TSSs of genes repressed in *Tdg*-null ECCs, they also overlapped in a similar proportion with TSSs of activated genes (data not shown). This argues against a simple model in which the co-engagement of TDG at DNMT3b-bound TSSs would prevent DNA methylation by inhibiting DNMT3B activity, thus favoring gene activity. Instead, we propose that an active methylation/demethylation reaction cycle is required to enable positive gene regulation. Consistent with this model, previous mapping of 5fC and 5caC through MAB-seq indicated that a fraction of these oxidized bases is found at promoters, including in CpG islands, although the role of TDG at promoters remained elusive in this case (Neri et al., 2015). Nonetheless, highly expressed gene promoters appear to undergo DNA methylation cycling that would be disrupted in *Tdg*-null cells. The nucleosome maps we have generated suggest that TDG fine-tunes nucleosome positioning at promoters as well as at other genomic sites such as CTCF binding sites. Indeed, the - 1/+1 nucleosomes around TSSs and those immediately flanking CTCF binding sites were differently positioned in the absence of TDG, consistent with a direct impact of 5mC turnover on the accurate positioning of specific nucleosomes. DNA methylation has already been proposed to favor nucleosome positioning and stability (Chodavarapu et al., 2010) but other studies reported opposite relationship (Li et al., 2022). In addition, 5fC has been shown to associate with well-positioned nucleosomes, likely through its capacity to directly interact with histones (Teif et al., 2014, Raiber et al., 2018), and sites that gain 5mC in Tet1/Tet2 KO ESCs also have a higher nucleosome occupancy (Wiehle et al., 2019). Hence, part of the transcriptional effect of TDG depletion may very well arise from alterations in nucleosome positioning caused by defects in 5mC turnover. Accordingly, we observed that nucleosomes flanking CTCF sites are shifted 30 bp towards CTCF sites in *Tdg*-null ECCs compared to wt ECCs, whereas +2, +3 and +4 nucleosome positions remain unchanged. The size of this nucleosome-shift effect is very reminiscent of that observed upon alterations in the activity of chromatin remodelers such as BRG1 (Hu et al., 2011; Ren et al., 2024). Hence, TDG could participate in coordinating nucleosome positioning and gene expression through functional interaction with chromatin remodelers such as CHD4, which associates with TDG binding sites (this study), and also with CTCF binding sites (Clarkson et al., 2019) and active promoters, where it participates to the definition of the NDR in mouse ESCs (de Dieuleveult et al., 2016). Although moderate, the estimated 40 bp narrowing of the NDR in *Tdg*-null cells is highly comparable to the 33 bp shrinkage of the NDR observed in yeast cells depleted of the chromatin remodeler RSC (Ganguli et al., 2014). Therefore, we propose that upon gene activation, TDG molecules that are pre-bound or newly recruited at promoters participate in CpG methylation/demethylation cycling and interact functionally with chromatin remodelers, which together allow for a proper positioning of the nucleosomes flanking the NDR, thereby facilitating access of NDR-binding transcription factors such as ATF4 to promoters.

In this study we also evidenced a functional link between TDG and ATF4-dependent transcription of ISR genes. Indeed, TDG binds to the promoters of *Atf4* and ATF4-target genes in ECCs, and a large subset of these ATF4-target genes are down-regulated in *Tdg*-null cells. This global effect likely results from the integration of several interdependent molecular mechanisms that include: regulation of ATF4 and its partner C/EBPb binding to DNA by 5mC turnover, nucleosome (re)positioning and acetylation of H3K27, and control of ATF4 protein levels via regulating mRNA stability and translation of its mRNA (Fig. 5K - Wortel et al., 2017). DNA binding of ATF4 and C/EBPb occurs at 5’-TGACGTCA-3’ sequences and ATF4 binding is inhibited by methylation of the central CpG (Kribelbauer et al., 2017; Yin et al, 2017). ATF4-C/EBPb heterodimers also bind to the non-canonical sequence 5’-CGATGCAA-3’ with higher affinity when the CpG is methylated (Mann et al., 2013). In addition, C/EBPb binds with a slightly higher affinity to the 5’-TTGCGTCA-3’ motif when the CpG is methylated, formylated or carboxylated (Sayeed et al., 2015; Yin et al., 2017). Given the documented impact of CpG methylation status on ATF4 and C/EBPb binding to DNA, a complete ablation of TDG activity is predicted to result in altered chromatin binding of these major stress integrators in various ways depending on the sequence of the binding site and the type of dimer engaged.

Upon stress conditions like aa starvation, ATF4 protein levels rise, favoring transcription of ATF4-target genes, the products of which act in a concerted manner during ISR. Members of the ISR pathway include aa synthetases, tRNA charging and solute carrier genes (Wortel et al., 2017; Lines et al., 2023). Here we show that both RA-induced cell aggregation and the mere formation of embryoid bodies are associated with a decrease in several *Slc* mRNAs encoding amino-acid transporters, including the leucine transporter SLC7A5. This suggests that aggregate formation induces leucine shortage, hence lowering mTORC1 activity through sestrin activation, and thereby reducing ATF4-dependent gene expression. In wt ECCs, treatment with RA leads to a TDG-dependent down-regulation of the leucine sensor and mTORC1 inhibitor *Sesn1*, thus likely increasing ATF4 production to sustain ATF4-target gene expression. Importantly, the implication of TDG in down-regulating *Sesn1* in ECCs is in line with the inverse correlation between *TDG* and *SESN1* expression in cancer patients. The cellular stress caused by tridimensional growth, which is reflected by the down-regulation of *Slc* genes, was found to be amplified in *Tdg*-null cells, implying a defective stress response upon *Tdg* inactivation. Amino-acid levels are sensed by two major pathways involving GCN2- and mTOR-regulated translation of ATF4 mRNA (Harding et al., 2000; Park et al., 2017; Torrence et al., 2021). In addition, mTORC1 induces a stabilization of ATF4 mRNA (Park et al., 2017). In the GCN2-ATF4 pathway, amino-acid starvation results in an increase in uncharged tRNAs, which bind to and activate the kinase GCN2, leading to phosphorylation of eIF2A and selective translation of the ATF4 mRNA. In growth factor-stimulated cells, uptake of essential amino-acids through SLC7A5 activates mTORC1, which allows for sustained growth and inhibition of TFE3-regulated autophagy (Nicklin et al., 2009). Our data unequivocally identified the mTORC1 pathway as being required for ATF4 activity in P19 ECCs and being affected in *Tdg*-null cells. In particular, depletion of TDG down-regulates *Slc7a5* mRNA levels in RA-treated cells. Also, the down-regulation of *Slc7a11* and *Slc3a2* in *Tdg*-null cells might prime cells for ferroptosis and participate in the reduction of mTORC1 activity (Zhang et al., 2021). In line with our observations, the mTORC1 pathway is involved in pluripotency exit of ESCs through TFE3 inhibition (Betschinger et al., 2013; Villegas et al., 2019). Furthermore, and consistent with the differentiation skewing observed in the absence of TDG, mTORC1 inhibition favors differentiation of ESCs into cardiomyocytes (Zheng et al., 2017). Thus, by controlling the activity of mTORC1 and ATF4, TDG takes part in metabolic reprogramming during cell differentiation, a role that may also well be of relevance to tumor cell growth and survival in human cancers.

## Acknowledgements

We thank L. Deleurme and A. Aimé for conducting cell sorting experiments at the Biosit cell sorting platform (https://biosit.univ-rennes.fr/cytometrie-en-flux-et-tri-cellulaire). We also thank V. Dupé for providing the Pax6 antibody, M. Pucéat for providing the Acta2 antibody, E. Chevet and E. Lafont for the gift of p70S6K and Phospho-p70S6K antibodies, PA Bidaud-Meynard for the gift of rapamycin, and A. Sérandour for providing sequencing reagents. We are grateful to R. Gibeaux for access to the fluorescence microscope and to A. Laurent for MNase-seq library preparation. Appreciation is extended to the Genomics Core Facility GenoA and the Bioinformatics Core Facility BiRD, both members of Biogenouest and France Genomique, as well as the Institut Français de Bioinformatique for their resources and technical support.

## Author contributions

MT, TM, EW, RA and MB: methodology and investigation. GB: methodology and formal analysis. SA: software and data curation. KS, GP, CF, MB, JP and CLP: investigation. GS: conceptualization, project administration, funding acquisition, investigation, visualization and original draft preparation. All authors reviewed and edited the manuscript.

## References

Abril-Garrido J, Dienemann C, Grabbe F, Velychko T, Lidschreiber M, Wang H, Cramer P. (2023). Structural basis of transcription reduction by a promoter-proximal +1 nucleosome. Mol Cell, 83(11):1798–1809.e7.

Ahola S, Rivera Mejías P, Hermans S, Chandragiri S, Giavalisco P, Nolte H, Langer T. (2022). OMA1-mediated integrated stress response protects against ferroptosis in mitochondrial cardiomyopathy. Cell Metab, 34(11):1875–1891.e7.

Aranda S, Alcaine-Colet A, Ballaré C, Blanco E, Mocavini I, Sparavier A, Vizán P, Borràs E, Sabidó E, Di Croce L. (2023). Thymine DNA glycosylase regulates cell-cycle-driven p53 transcriptional control in pluripotent cells. Mol Cell, 83(15):2673–2691.e7.

Bai L, Morozov AV. (2010). Gene regulation by nucleosome positioning. Trends Genet, 26(11):476–83.

Betschinger J, Nichols J, Dietmann S, Corrin PD, Paddison PJ, Smith A. (2013). Exit from pluripotency is gated by intracellular redistribution of the bHLH transcription factor Tfe3. Cell, 153(2):335–47.

Bian Z, Sun X, Liu L, Qin Y, Zhang Q, Liu H, Mao L, Sun S. (2023). Sodium Butyrate Induces CRC Cell Ferroptosis via the CD44/SLC7A11 Pathway and Exhibits a Synergistic Therapeutic Effect with Erastin. Cancers (Basel),15(2):423.

Boland MJ, Christman JK. (2008). Characterization of Dnmt3b:thymine-DNA glycosylase interaction and stimulation of thymine glycosylase-mediated repair by DNA methyltransferase(s) and RNA. J Mol Biol, 379(3):492–504.

Cangelosi AL, Puszynska AM, Roberts JM, Armani A, Nguyen TP, Spinelli JB, Kunchok T, Wang B, Chan SH, Lewis CA, Comb WC, Bell GW, Helman A, Sabatini DM. (2022). Zonated leucine sensing by Sestrin-mTORC1 in the liver controls the response to dietary leucine. Science, 377(6601):47–56.

Caron G, Hussein M, Kulis M, Delaloy C, Chatonnet F, Pignarre A, Avner S, Lemarié M, Mahé EA, Verdaguer-Dot N, Queirós AC, Tarte K, Martín-Subero JI, Salbert G, Fest T. (2015). Cell-Cycle-Dependent Reconfiguration of the DNA Methylome during Terminal Differentiation of Human B Cells into Plasma Cells. Cell Rep, 13(5):1059–71.

Carroll T, Barrows D. (2024). profileplyr: Visualization and annotation of read signal over genomic ranges with profileplyr. R package version 1.22.0.

Charpentier E, Cornec M, Dumont S, Meistermann D, Bordron P, David L, Redon R, Bonnaud S, Bihouée A. (2021). 3’ RNA sequencing for rapid and low cost gene expression profiling. Protocolexchange, 10.21203/rs.3.pex-1336/v1.

Chaumette T, Cinotti R, Mollé A, Solomon P, Castain L, Fourgeux C, McWilliam HEG, Misme-Aucouturier B, Broquet A, Jacqueline C, Vourc’h M, Fradin D, Bossard C, David L, Montassier E, Braudeau C, Josien R, Villadangos JA, Asehnoune K, Bressollette-Bodin C, Poschmann J, Roquilly A. (2022). Monocyte Signature Associated with Herpes Simplex Virus Reactivation and Neurological Recovery after Brain Injury. Am J Respir Crit Care Med, 206(3):295–310.

Cheng S, Mittnenzweig M, Mayshar Y, Lifshitz A, Dunjić M, Rais Y, Ben-Yair R, Gehrs S, Chomsky E, Mukamel Z, Rubinstein H, Schlereth K, Reines N, Orenbuch AH, Tanay A, Stelzer Y. (2022). The intrinsic and extrinsic effects of TET proteins during gastrulation. Cell, 185(17):3169–3185.e20.

Chodavarapu RK, Feng S, Bernatavichute YV, Chen PY, Stroud H, Yu Y, Hetzel JA, Kuo F, Kim J, Cokus SJ, Casero D, Bernal M, Huijser P, Clark AT, Krämer U, Merchant SS, Zhang X, Jacobsen SE, Pellegrini M. (2010). Relationship between nucleosome positioning and DNA methylation. Nature, 466(7304):388–92.

Clarkson CT, Deeks EA, Samarista R, Mamayusupova H, Zhurkin VB, Teif VB. (2019). CTCF-dependent chromatin boundaries formed by asymmetric nucleosome arrays with decreased linker length. Nucleic Acids Res, 47(21):11181–11196.

Cortázar D, Kunz C, Selfridge J, Lettieri T, Saito Y, MacDougall E, Wirz A, Schuermann D, Jacobs AL, Siegrist F, Steinacher R, Jiricny J, Bird A, Schär P. (2011). Embryonic lethal phenotype reveals a function of TDG in maintaining epigenetic stability. Nature, 470(7334):419–23.

D’Aniello C, Fico A, Casalino L, Guardiola O, Di Napoli G, Cermola F, De Cesare D, Tatè R, Cobellis G, Patriarca EJ, Minchiotti G. (2015). A novel autoregulatory loop between the Gcn2-Atf4 pathway and (L)-Proline [corrected] metabolism controls stem cell identity. Cell Death Differ, 22(7):1094–105.

Deaton AM, Bird A. (2011). CpG islands and the regulation of transcription. Genes Dev, 25(10):1010–22.

de Dieuleveult M, Yen K, Hmitou I, Depaux A, Boussouar F, Bou Dargham D, Jounier S, Humbertclaude H, Ribierre F, Baulard C, Farrell NP, Park B, Keime C, Carrière L, Berlivet S, Gut M, Gut I, Werner M, Deleuze JF, Olaso R, Aude JC, Chantalat S, Pugh BF, Gérard M. (2016). Genome-wide nucleosome specificity and function of chromatin remodellers in ES cells. Nature, 530(7588):113–6.

Dixon SJ, Lemberg KM, Lamprecht MR, Skouta R, Zaitsev EM, Gleason CE, Patel DN, Bauer AJ, Cantley AM, Yang WS, Morrison B 3rd, Stockwell BR. (2012). Ferroptosis: an iron-dependent form of nonapoptotic cell death. Cell, 149(5):1060-72.

Edri S, Hayward P, Baillie-Johnson P, Steventon BJ, Martinez Arias A. (2019). An epiblast stem cell-derived multipotent progenitor population for axial extension. Development, 146(10):dev168187.

ENCODE Project Consortium, Moore JE, Purcaro MJ, Pratt HE, Epstein CB, Shoresh N, Adrian J, et al. (2020). Expanded Encyclopaedias of DNA Elements in the Human and Mouse Genomes. Nature, 583(7818):699–710.

Faletti S, Osti D, Ceccacci E, Richichi C, Costanza B, Nicosia L, Noberini R, Marotta G, Furia L, Faretta MR, Brambillasca S, Quarto M, Bertero L, Boldorini R, Pollo B, Gandini S, Cora D, Minucci S, Mercurio C, Varasi M, Bonaldi T, Pelicci G. (2021). LSD1-directed therapy affects glioblastoma tumorigenicity by deregulating the protective ATF4-dependent integrated stress response. Sci Transl Med, 13(623):eabf7036.

Gallais R, Demay F, Barath P, Finot L, Jurkowska R, Le Guével R, Gay F, Jeltsch A, Métiver R, Salbert G. (2007) Deoxyribonucleic Acid Methyl Transferases 3a and 3b Associate with the Nuclear Orphan Receptor COUP-TFI during Gene Activation. Mol Endocrinol, 21:2085–2096.

Ganguli D, Chereji RV, Iben JR, Cole HA, Clark DJ. (2014). RSC-dependent constructive and destructive interference between opposing arrays of phased nucleosomes in yeast. Genome Res, 24(10):1637–49.

Ginno PA, Gaidatzis D, Feldmann A, Hoerner L, Imanci D, Burger L, Zilbermann F, Peters AHFM, Edenhofer F, Smallwood SA, Krebs AR, Schübeler D. (2020). A genome-scale map of DNA methylation turnover identifies site-specific dependencies of DNMT and TET activity. Nat Commun, 11(1):2680.

Han DW, Tapia N, Araúzo-Bravo MJ, Lim KT, Kim KP, Ko K, Lee HT, Schöler HR. (2013). Sox2 Level Is a Determinant of Cellular Reprogramming Potential. PLoS One, 8(6):e67594.

Harding HP, Novoa I, Zhang Y, Zeng H, Wek R, Schapira M, Ron D. (2000). Regulated translation initiation controls stress-induced gene expression in mammalian cells. Mol Cell, 6(5):1099–108.

He F, Zhang P, Liu J, Wang R, Kaufman RJ, Yaden BC, Karin M. (2023). ATF4 suppresses hepatocarcinogenesis by inducing SLC7A11 (xCT) to block stress-related ferroptosis. J Hepatol, 79(2):362–377.

Henrique D, Abranches E, Verrier L, Storey KG. (2015). Neuromesodermal progenitors and the making of the spinal cord. Development, 142(17):2864–75.

Hu G, Schones DE, Cui K, Ybarra R, Northrup D, Tang Q, Gattinoni L, Restifo NP, Huang S, Zhao K. (2011). Regulation of nucleosome landscape and transcription factor targeting at tissue-specific enhancers by BRG1. Genome Res, 21(10):1650–8.

Jiang J, Srivastava S, Seim G, Pavlova NN, King B, Zou L, Zhang C, Zhong M, Feng H, Kapur R, Wek RC, Fan J, Zhang J. (2019). Promoter demethylation of the asparagine synthetase gene is required for ATF4-dependent adaptation to asparagine depletion. J Biol Chem, 294:18674–84.

Jiang X, Stockwell BR, Conrad M. (2021). Ferroptosis: mechanisms, biology and role in disease. Nat Rev Mol Cell Biol, 22(4):266–282.

Jones-Villeneuve EM, McBurney MW, Rogers KA, Kalnins VI. (1982). Retinoic acid induces embryonal carcinoma cells to differentiate into neurons and glial cells. J Cell Biol, 94(2):253–62.

Kaluscha S, Domcke S, Wirbelauer C, Stadler MB, Durdu S, Burger L, Schübeler D. (2022). Evidence that direct inhibition of transcription factor binding is the prevailing mode of gene and repeat repression by DNA methylation. Nat Genet, 54(12):1895–1906.

Kim HB, Choi WY, Kim SH, Lee HJ, Park SG. (2024). GNAO1: A Novel Tumor Suppressor Gene in Colorectal Cancer Pathogenesis. Anticancer Res, 44(12):5425–5433.

Kriaucionis S, Heintz N. (2009). The nuclear DNA base 5-hydroxymethylcytosine is present in Purkinje neurons and the brain. Science, 324(5929):929–30.

Kribelbauer JF, Laptenko O, Chen S, Martini GD, Freed-Pastor WA, Prives C, Mann RS, Bussemaker HJ. (2017). Quantitative Analysis of the DNA Methylation Sensitivity of Transcription Factor Complexes. Cell Rep, 19(11):2383–2395.

Kuang F, Liu J, Li C, Kang R, Tang D. (2020). Cathepsin B is a mediator of organelle-specific initiation of ferroptosis. Biochem Biophys Res Commun, 533(4):1464–1469.

Kuang F, Liu J, Xie Y, Tang D, Kang R. (2021). MGST1 is a redox-sensitive repressor of ferroptosis in pancreatic cancer cells. Cell Chem Biol, 28(6):765–775.e5.

Lai K, Song C, Gao M, Deng Y, Lu Z, Li N, Geng Q. (2023). Uridine Alleviates Sepsis-Induced Acute Lung Injury by Inhibiting Ferroptosis of Macrophage. Int J Mol Sci, 24(6):5093.

Langmead B, Trapnell C, Pop M, Salzberg SL. (2009). Ultrafast and memory-efficient alignment of short DNA sequences to the human genome. Genome Biol, 10(3):R25.

Langton S, Gudas LJ. (2008). CYP26A1 knockout embryonic stem cells exhibit reduced differentiation and growth arrest in response to retinoic acid. Dev Biol, 315(2):331–54.

Lassot I, Estrabaud E, Emiliani S, Benkirane M, Benarous R, Margottin-Goguet F. (2005). p300 modulates ATF4 stability and transcriptional activity independently of its acetyltransferase domain. J Biol Chem, 280(50):41537–45.

Laurent A, Madigou T, Bizot M, Turpin M, Palierne G, Mahé E, Guimard S, Métivier R, Avner S, Le Péron C, Salbert G. (2022). TET2-mediated epigenetic reprogramming of breast cancer cells impairs lysosome biogenesis. Life Sci Alliance, 5(7):e202101283.

Lauria A, Meng G, Proserpio V, Rapelli S, Maldotti M, Polignano IL, Anselmi F, Incarnato D, Krepelova A, Donna D, Levra Levron C, Donati G, Molineris I, Neri F, Oliviero S. (2023). DNMT3B supports meso-endoderm differentiation from mouse embryonic stem cells. Nat Commun, 14(1):367.

Letellier T, Kervella D, Sadek A, Masset C, Garandeau C, Fourgeux C, Gourain V, Poschmann J, Blancho G, Ville S, On Behalf of The Divat Consortium. (2022). Time-Limited Therapy with Belatacept in Kidney Transplant Recipients. J Clin Med, 11(11):3229.

Li YQ, Zhou PZ, Zheng XD, Walsh CP, Xu GL. (2007). Association of Dnmt3a and thymine DNA glycosylase links DNA methylation with base-excision repair. Nucleic Acids Res, 35(2):390–400.

Li H, Handsaker B, Wysoker A, Fennell T, Ruan J, Homer N, Marth G, Abecasis G, Durbin R; 1000 Genome Project Data Processing Subgroup. (2009). The Sequence Alignment/Map format and SAMtools. Bioinformatics, 25(16):2078-9.

Li X, Yue X, Pastor WA, Lin L, Georges R, Chavez L, Evans SM, Rao A. (2016). Tet proteins influence the balance between neuroectodermal and mesodermal fate choice by inhibiting Wnt signaling. Proc Natl Acad Sci U S A, 113(51):E8267–E8276.

Li Z, Feng Y, Zhang Z, Cao X, Lu X. (2020). TMPO-AS1 promotes cell proliferation of thyroid cancer via sponging miR-498 to modulate TMPO. Cancer Cell Int, 20:294.

Li S, Peng Y, Panchenko AR. (2022). DNA methylation: Precise modulation of chromatin structure and dynamics. Curr Opin Struct Biol, 75:102430.

Lines CL, McGrath MJ, Dorwart T, Conn CS. (2023). The integrated stress response in cancer progression: a force for plasticity and resistance. Front Oncol, 13:1206561.

Liu T, Ortiz JA, Taing L, Meyer CA, Lee B, Zhang Y, Shin H, Wong SS, Ma J, Lei Y, Pape UJ, Poidinger M, Chen Y, Yeung K, Brown M, Turpaz Y, Liu XS. (2011). Cistrome: an integrative platform for transcriptional regulation studies. Genome Biol, 12(8):R83.

Liu Q, Wang F, Chen Y, Cui H, Wu H. (2024). A regulatory module comprising G3BP1-FBXL5-IRP2 axis determines sodium arsenite-induced ferroptosis. J Hazard Mater, 465:133038.

Love MI, Huber W, Anders S. (2014). Moderated estimation of fold change and dispersion for RNA-seq data with DESeq2. Genome Biology, 15:550. 10.1186/s13059-014-0550-8

Magnúsdóttir E, Dietmann S, Murakami K, Günesdogan U, Tang F, Bao S, Diamanti E, Lao K, Gottgens B, Azim Surani M. (2013). A tripartite transcription factor network regulates primordial germ cell specification in mice. Nat Cell Biol, 15(8):905–15.

Mahé EA, Madigou T, Sérandour AA, Bizot M, Avner S, Chalmel F, Palierne G, Métivier R, Salbert G. (2017). Cytosine modifications modulate the chromatin architecture of transcriptional enhancers. Genome Res, 27(6):947–958.

Maiti A, Drohat AC. (2011). Thymine DNA glycosylase can rapidly excise 5-formylcytosine and 5-carboxylcytosine: potential implications for active demethylation of CpG sites. J Biol Chem, 286(41):35334–38.

Mann IK, Chatterjee R, Zhao J, He X, Weirauch MT, Hughes TR, Vinson C. (2013). CG methylated microarrays identify a novel methylated sequence bound by the CEBPB|ATF4 heterodimer that is active in vivo. Genome Res, 23(6):988–97.

McBurney MW. (1993). P19 embryonal carcinoma cells. Int J Dev Biol, 37(1):135–40.

Ménoret S, Tesson L, Remy S, Gourain V, Sérazin C, Usal C, Guiffes A, Chenouard V, Ouisse LH, Gantier M, Heslan JM, Fourgeux C, Poschmann J, Guillonneau C, Anegon I. (2023). CD4^+^ and CD8^+^ regulatory T cell characterization in the rat using a unique transgenic Foxp3-EGFP model. BMC Biol, 21(1):8.

Métivier R, Gallais R, Tiffoche C, Le Péron C, Jurkowska RZ, Carmouche RP, Ibberson D, Barath P, Demay F, Reid G, Benes V, Jeltsch A, Gannon F, Salbert G. (2008). Cyclical DNA methylation of a transcriptionally active promoter. Nature, 452(7183):45–50.

Mulholland CB, Nishiyama A, Ryan J, Nakamura R, Yiğit M, Glück IM, Trummer C, Qin W, Bartoschek MD, Traube FR, Parsa E, Ugur E, Modic M, Acharya A, Stolz P, Ziegenhain C, Wierer M, Enard W, Carell T, Lamb DC, Takeda H, Nakanishi M, Bultmann S, Leonhardt H. (2020). Recent evolution of a TET-controlled and DPPA3/STELLA-driven pathway of passive DNA demethylation in mammals. Nat Commun, 11(1):5972.

Nan X, Ng HH, Johnson CA, Laherty CD, Turner BM, Eisenman RN, Bird A. (1998). Transcriptional repression by the methyl-CpG-binding protein MeCP2 involves a histone deacetylase complex. Nature, 393(6683):386–9.

Neddermann P, Jiricny J. (1993). The purification of a mismatch-specific thymine-DNA glycosylase from HeLa cells. J Biol Chem, 268(28):21218–24.

Neddermann P, Jiricny J. (1994). Efficient removal of uracil from G.U mispairs by the mismatch-specific thymine DNA glycosylase from HeLa cells. Proc Natl Acad Sci U S A, 91(5):1642–6.

Neri F, Incarnato D, Krepelova A, Rapelli S, Anselmi F, Parlato C, Medana C, Dal Bello F, Oliviero S. (2015). Single-Base Resolution Analysis of 5-Formyl and 5-Carboxyl Cytosine Reveals Promoter DNA Methylation Dynamics. Cell Rep, 10(5):674–683.

Nicklin P, Bergman P, Zhang B, Triantafellow E, Wang H, Nyfeler B, Yang H, Hild M, Kung C, Wilson C, Myer VE, MacKeigan JP, Porter JA, Wang YK, Cantley LC, Finan PM, Murphy LO. (2009). Bidirectional transport of amino acids regulates mTOR and autophagy. Cell, 136(3):521–34.

Okada Y, Shimazaki T, Sobue G, Okano H. (2004). Retinoic-acid-concentration-dependent acquisition of neural cell identity during in vitro differentiation of mouse embryonic stem cells. Dev Biol, 275(1):124–42.

Onabote O, Hassan HM, Isovic M, Torchia J. (2022). The Role of Thymine DNA Glycosylase in Transcription, Active DNA Demethylation, and Cancer. Cancers (Basel), 14(3):765.

Pakos-Zebrucka K, Koryga I, Mnich K, Ljujic M, Samali A, Gorman AM. (2016). The integrated stress response. EMBO Rep, 17(10):1374–1395.

Parisi S, D’Andrea D, Lago CT, Adamson ED, Persico MG, Minchiotti G. (2003). Nodal-dependent Cripto signaling promotes cardiomyogenesis and redirects the neural fate of embryonic stem cells. J Cell Biol, 163(2):303–14.

Park Y, Reyna-Neyra A, Philippe L, Thoreen CC. (2017). mTORC1 Balances Cellular Amino Acid Supply with Demand for Protein Synthesis through Post-transcriptional Control of ATF4. Cell Rep, 19(6):1083–1090.

Parry A, Rulands S, Reik, W. (2021). Active turnover of DNA methylation during cell fate decisions. Nature reviews. Genetics, 22(1), 59–66. https://doi-org.insb.bib.cnrs.fr/10.1038/s41576-020-00287-8

Raiber EA, Portella G, Martínez Cuesta S, Hardisty R, Murat P, Li Z, Iurlaro M, Dean W, Spindel J, Beraldi D, Liu Z, Dawson MA, Reik W, Balasubramanian S. (2018). 5-Formylcytosine organizes nucleosomes and forms Schiff base interactions with histones in mouse embryonic stem cells. Nat Chem, 10(12):1258–1266.

Ramírez F, Ryan DP, Grüning B, Bhardwaj V, Kilpert F, Richter AS, Heyne S, Dündar F, Manke T. (2016). deepTools2: A next Generation Web Server for Deep-Sequencing Data Analysis. Nucleic Acids Research, 44(W1). doi:10.1093/nar/gkw257.

Ren G, Ku WL, Ge G, Hoffman JA, Kang JY, Tang Q, Cui K, He Y, Guan Y, Gao B, Liu C, Archer TK, Zhao K. (2024). Acute depletion of BRG1 reveals its primary function as an activator of transcription. Nat Commun, 15(1):4561.

Rulands S, Lee HJ, Clark SJ, Angermueller C, Smallwood SA, Krueger F, Mohammed H, Dean W, Nichols J, Rugg-Gunn P, Kelsey G, Stegle O, Simons BD, Reik W. (2018). Genome-Scale Oscillations in DNA Methylation during Exit from Pluripotency. Cell Syst, 7(1):63–76.e12.

Russo L, Sladitschek HL, Neveu PA. (2022). Multi-layered regulation of neuroectoderm differentiation by retinoic acid in a primitive streak-like context. Stem Cell Reports, 17(2):231–244.

Sandoval JE, Huang YH, Muise A, Goodell MA, Reich NO. (2019). Mutations in the DNMT3A DNA methyltransferase in acute myeloid leukemia patients cause both loss and gain of function and differential regulation by protein partners. J Biol Chem, 294(13):4898–4910.

Sayeed SK, Zhao J, Sathyanarayana BK, Golla JP, Vinson C. (2015). C/EBPβ (CEBPB) protein binding to the C/EBP|CRE DNA 8-mer TTGC|GTCA is inhibited by 5hmC and enhanced by 5mC, 5fC, and 5caC in the CG dinucleotide. Biochim Biophys Acta, 1849(6):583-9.

Sérandour AA, Avner S, Oger F, Bizot M, Percevault F, Lucchetti-Miganeh C, Palierne G, Gheeraert C, Barloy-Hubler F, Péron CL, Madigou T, Durand E, Froguel P, Staels B, Lefebvre P, Métivier R, Eeckhoute J, Salbert G. (2012). Dynamic hydroxymethylation of deoxyribonucleic acid marks differentiation-associated enhancers. Nucleic Acids Res, 40(17):8255–65.

Shao S, Liu Y, Hong W, Mo Y, Shu F, Jiang L, Tan N. (2023). Influence of FOSL1 Inhibition on Vascular Calcification and ROS Generation through Ferroptosis via P53-SLC7A11 Axis. Biomedicines, 11(2):635.

Steinacher R, Barekati Z, Botev P, Kuśnierczyk A, Slupphaug G, Schär P. (2019). SUMOylation coordinates BERosome assembly in active DNA demethylation during cell differentiation. EMBO J, 38(1):e99242.

Stephens M. (2016). False discovery rates: a new deal. Biostatistics, 18:2. 10.1093/biostatistics/kxw041

Swanda RV, Ji Q, Wu X, Yan J, Dong L, Mao Y, Uematsu S, Dong Y, Qian SB. (2023). Lysosomal cystine governs ferroptosis sensitivity in cancer via cysteine stress response. Mol Cell, 83(18):3347–3359.e9.

Tahiliani M, Koh KP, Shen Y, Pastor WA, Bandukwala H, Brudno Y, Agarwal S, Iyer LM, Liu DR, Aravind L, Rao A. (2009). Conversion of 5-methylcytosine to 5-hydroxymethylcytosine in mammalian DNA by MLL partner TET1. Science, 324(5929):930–5.

Teif VB, Beshnova DA, Vainshtein Y, Marth C, Mallm JP, Höfer T, Rippe K. (2014). Nucleosome repositioning links DNA (de)methylation and differential CTCF binding during stem cell development. Genome Res, 24(8):1285–95.

Tian X, Zhang S, Zhou L, Seyhan AA, Hernandez Borrero L, Zhang Y, El-Deiry WS. (2021). Targeting the Integrated Stress Response in Cancer Therapy. Front Pharmacol, 12:747837.

Tini M, Benecke A, Um SJ, Torchia J, Evans RM, Chambon P. (2002). Association of CBP/p300 acetylase and thymine DNA glycosylase links DNA repair and transcription. Mol Cell, 9(2):265–77.

Torrence ME, MacArthur MR, Hosios AM, Valvezan AJ, Asara JM, Mitchell JR, Manning BD. (2021). The mTORC1-mediated activation of ATF4 promotes protein and glutathione synthesis downstream of growth signals. Elife, 10:e63326.

Um S, Harbers M, Benecke A, Pierrat B, Losson R, Chambon P. (1998). Retinoic acid receptors interact physically and functionally with the T:G mismatch-specific thymine-DNA glycosylase. J Biol Chem, 273(33):20728–36.

Untergasser A, Cutcutache I, Koressaar T, Ye J, Faircloth BC, Remm M and Rozen SG. (2012). Primer3--new capabilities and interfaces. Nucleic Acids Res, 40(15):e115.

Villegas F, Lehalle D, Mayer D, Rittirsch M, Stadler MB, Zinner M, Olivieri D, Vabres P, Duplomb-Jego L, De Bont ESJM, Duffourd Y, Duijkers F, Avila M, Geneviève D, Houcinat N, Jouan T, Kuentz P, Lichtenbelt KD, Thauvin-Robinet C, St-Onge J, Thevenon J, van Gassen KLI, van Haelst M, van Koningsbruggen S, Hess D, Smallwood SA, Rivière JB, Faivre L, Betschinger J. (2019). Lysosomal Signaling Licenses Embryonic Stem Cell Differentiation via Inactivation of Tfe3. Cell Stem Cell, 24(2):257–270.e8.

Voong LN, Xi L, Sebeson AC, Xiong B, Wang JP, Wang X. (2016). Insights into Nucleosome Organization in Mouse Embryonic Stem Cells through Chemical Mapping. Cell, 167(6):1555–1570.e15.

Watrin E, Demidova M, Watrin T, Hu Z, Prigent C. (2014). Sororin pre-mRNA splicing is required for proper sister chromatid cohesion in human cells. EMBO Rep, 15(9):948–55.

Wiehle L, Thorn GJ, Raddatz G, Clarkson CT, Rippe K, Lyko F, Breiling A, Teif VB. (2019). DNA (de)methylation in embryonic stem cells controls CTCF-dependent chromatin boundaries. Genome Res, 29(5):750–761.

Wheldon LM, Abakir A, Ferjentsik Z, Dudnakova T, Strohbuecker S, Christie D, Dai N, Guan S, Foster JM, Corrêa IR Jr, Loose M, Dixon JE, Sottile V, Johnson AD, Ruzov A. (2014). Transient accumulation of 5-carboxylcytosine indicates involvement of active demethylation in lineage specification of neural stem cells. Cell Rep, 7(5):1353–1361.

Wickham H. (2016). ggplot2: Elegant Graphics for Data Analysis. Springer-Verlag New York. ISBN 978-3-319-24277-4, https://ggplot2.tidyverse.org.

Wolfson RL, Chantranupong L, Saxton RA, Shen K, Scaria SM, Cantor JR, Sabatini DM. (2016). Sestrin2 is a leucine sensor for the mTORC1 pathway. Science, 351(6268):43–8.

Wortel IMN, van der Meer LT, Kilberg MS, van Leeuwen FN. (2017). Surviving Stress: Modulation of ATF4-Mediated Stress Responses in Normal and Malignant Cells. Trends Endocrinol Metab, 28(11):794–806.

Wu X, Zhang Y. (2017). TET-mediated active DNA demethylation: mechanism, function and beyond. Nat Rev Genet, 18(9):517–534.

Xia J, Fu B, Wang Z, Wen G, Gu Q, Chen D, Ren H. (2024). MVP enhances FGF21-induced ferroptosis in hepatocellular carcinoma by increasing lipid peroxidation through regulation of NOX4. Clin Transl Sci, 17(8):e13910.

Yang WS, SriRamaratnam R, Welsch ME, Shimada K, Skouta R, Viswanathan VS, Cheah JH, Clemons PA, Shamji AF, Clish CB, Brown LM, Girotti AW, Cornish VW, Schreiber SL, Stockwell BR. (2014). Regulation of ferroptotic cancer cell death by GPX4. Cell, 156(1-2):317–331.

Yang W, Mu B, You J, Tian C, Bin H, Xu Z, Zhang L, Ma R, Wu M, Zhang G, Huang C, Li L, Shao Z, Dai L, Désaubry L, Yang S. (2022). Non-classical ferroptosis inhibition by a small molecule targeting PHB2. Nat Commun, 13(1):7473.

Yao J, Chen X, Liu X, Li R, Zhou X, Qu Y. (2021). Characterization of a ferroptosis and iron-metabolism related lncRNA signature in lung adenocarcinoma. Cancer Cell Int, 21(1):340.

Yin Y, Morgunova E, Jolma A, Kaasinen E, Sahu B, Khund-Sayeed S, Das PK, Kivioja T, Dave K, Zhong F, Nitta KR, Taipale M, Popov A, Ginno PA, Domcke S, Yan J, Schübeler D, Vinson C, Taipale J. (2017). Impact of cytosine methylation on DNA binding specificities of human transcription factors. Science, 356(6337):eaaj2239.

Zhang Y, Liu T, Meyer CA, Eeckhoute J, Johnson DS, Bernstein BE, Nusbaum C, Myers RM, Brown M, Li W, Liu XS. (2008). Model-based analysis of ChIP-Seq (MACS). Genome Biol,; 9(9):R137.

Zhang Y, Swanda RV, Nie L, Liu X, Wang C, Lee H, Lei G, Mao C, Koppula P, Cheng W, Zhang J, Xiao Z, Zhuang L, Fang B, Chen J, Qian SB, Gan B. (2021). mTORC1 couples cyst(e)ine availability with GPX4 protein synthesis and ferroptosis regulation. Nat Commun, 12(1):1589.

Zheng B, Wang J, Tang L, Shi J, Zhu D. (2017). mTORC1 and mTORC2 play different roles in regulating cardiomyocyte differentiation from embryonic stem cells. Int J Dev Biol, 61(1-2):65–72.

Zhou Y, Wu H, Wang F, Xu L, Yan Y, Tong X, Yan H. (2022). GPX7 Is Targeted by miR-29b and GPX7 Knockdown Enhances Ferroptosis Induced by Erastin in Glioma. Front Oncol, 11:802124.

