## Supplementary Material for "TDG orchestrates ATF4-dependent gene transcription during retinoic acid-induced cell fate acquisition"

**Content:**

**Supplementary Figure Legends**

**Supplementary Figures**

**Supplementary Table 1**: List of oligonucleotides used in this study.

**Supplementary Table 2**: Gene expression fold change (Log2) and associated P values comparing untreated monolayers of *Tdg*-null ECCs and wt ECCs.

**Supplementary Table 3**: Gene expression fold change (Log2) and associated P values comparing RA-treated monolayers of *Tdg*-null ECCs and wt ECCs.

**Supplementary Figure Legends:**

**Supplementary Figure 1: Inactivation of *Tdg* in P19 ECCs.** (**A**) Time-course analysis of cell differentiation markers upon RA treatment. Normalized 3'RNA-seq data are expressed as counts per million reads (CPM). (**B**) PCR analysis of genomic DNA from various cell clones isolated after CRISPR/Cas9 mediated knock-out of the *Tdg* gene (left panel). The G2.10 clone underwent the expected recombination event and does not express the *Tdg* mRNA (lower lane: RT-PCR). RT-qPCR (middle panel) confirmed the absence of *Tdg* mRNA in clone G2.10. No TDG protein was detected by western blot (right panel). Arrowheads indicate TDG bands and the asterisk indicates a non-specific band. Meis1 detection serves as a positive control for RA treatment and U2AF65 as a loading control. (**C**) MA plot showing differentially expressed genes in monolayers of untreated *Tdg*-null ECCs compared to untreated wt ECCs (left panel). Up-regulated genes are highlighted in red and down-regulated genes in blue. Names of highly significantly differentially regulated genes are indicated. Gene ontology (GO) enrichment analysis of up- and down-regulated genes is shown on the right. (**D**) RT-qPCR assay of *Cyp26a1* mRNA levels in wt and *Tdg*-null cells treated with RA for the indicated times. (**E**) RT-qPCR assay of *Meis1* mRNA levels in wt and *Tdg*-null ECCs treated with increasing concentrations of RA or with 1% DMSO for 48 hours. (**F**) Phase-contrast microscopy images of wt and *Tdg*-null ECCs grown as monolayers and treated with increasing concentrations of RA or with 1% DMSO for 48 hours. White triangles point to regions of RA-induced 3D growth. Scale bar: 10 μm. (**G**) PCR analysis of genomic DNA from various cell clones isolated after CRISPR/Cas9 mediated knock-out of the *Cyp26a1* gene. Homozygote KO clones 11 and 16 were selected for further experiments. (**H**) RT-qPCR assay of *Cyp26a1* mRNA levels in the indicated cells treated (+RA) or not (-RA) with 1 μM RA for 48 hours.

**Supplementary Figure 2: TDG protects from cell death.** (**A**) Results of MTT assays in wt and *Tdg*-null ECCs treated with increasing concentrations of RA or with 1% DMSO. (**B**) Phase-contrast microscopy images of wt and *Tdg*-null ECCs grown for 48 hours with 1 μM RA. Images were acquired after one wash with PBS. Asterisks indicate dead floating cells. Scale bar: 30 μm. (**C**) List of 15 genes involved in ferroptosis regulation and belonging to the top 47 differentially expressed genes between *Tdg*-null and wt ECCs. For each gene, its positive (+) or negative (-) influence on ferroptosis as well as the Log2 fold change (*Tdg*-null versus wt ECCs) and the associated adjusted P value are indicated. (**D**) Phase-contrast microscopy images of wt and *Tdg*-null ECCs grown for 24 hours with 100 nM RA +/- 20 μM Fer1. Scale bar: 30 μm.

**Supplementary Figure 3: TDG sustains ATF4-dependent gene expression.** (**A**) Time course analysis of 1 μM RA effect on mRNA levels of the indicated genes determined by RT-qPCR. *Atf4*, *Asns*, *Chac1* and *Psat1* are ATF4-target genes, while *Nanog* and *Meis1* are used as pluripotency and differentiation markers respectively. (**B**) Time course study of 1 μM RA effect on mRNA levels for selected solute carrier genes (*Slc*) extracted from 3'RNA-seq data. (**C**) Integrated genome browser (IGB) snapshots of TDG and ATF4 ChIP-seq signal at *Slc* genes. (**D**) Model depicting how cell culture conditions influence the expression of ATF4-target genes. (**E**) RT-qPCR assays of *Asns* and *Psat1* mRNA levels in wt and *Tdg*-null ECCs treated for 48 hours with halofuginone (Halo) or L-proline (L-Pro). (**F**) Differential expression (RNA-seq) of genes involved either in activation or in inhibition of mTORC1, in wt and *Tdg*-null ECCs cultured in the indicated conditions. Significant differential expression levels are highlighted in green.

**Supplementary Figure 4: TDG impacts on nucleosome organization at TSSs and CTCF binding sites.** (**A**) Integrated genome browser (IGB) snapshots of TDG, ATF4 and CTCF ChIP-seq signal at selected ATF4-target genes. (**B**) Venn diagram displaying the overlaps between ESC and ECC CTCF binding sites (left panel). The right panel shows IGB snapshots of ESC and ECC CTCF ChIP-seq signal at selected loci. (**C**) Average profiles of MNase-seq signal at C3, C4 and C5 TSSs in monolayers of wt or *Tdg*-null ECCs treated or not with 1 μM RA for 48 hours. (**D**) Expression values (as Log of the mean of RNA-seq triplicates) of genes which TSSs are included in clusters C3, C4 and C5 from Fig. 4D in untreated wt and *Tdg*-null ECCs. (**E**) MNase-qPCR analysis of +1 nucleosome density at the indicated genes in wt or *Tdg*-null ECCs treated or not with 1 μM RA. MNase-qPCR values are reported to the no MNase (0 U) condition. (**F**) Average profiles of MNase-seq signal around C4, and C5 TSSs in wt or *Tdg*-null ECCs treated with 1 μM RA. The ATF4 ChIP-seq trace is also shown for each cluster. (**G**) Average profiles of MNase-seq signal at non-promoter CTCF binding sites common between ESCs and ECCs in wt and *Tdg*-null ECCs either untreated (left panel) or treated with 1 μM RA (right panel).

**Supplementary Figure 5: Relationship between TDG and ferroptosis genes in cancer.** (**A**) ) Heatmaps showing *TDG*, *TMPO* and *MVP* expression levels in lung tumor samples (TCGA lung cancer cohort, n=1,129) ranked according to *TDG* expression levels. (**B**) *FBXL5* and *GNAO1* gene effect in CRISPR experiments in cell lines with *TDG* damaging mutations (1) and cell lines with no identified mutations (0). The names of *TDG* mutant cell lines with an absolute gene effect ≥ 0.2 are shown. The distribution of *FBXL5* and *GNAO1* gene effects in all cell lines has been added for both CRISPR and RNAi experiments.

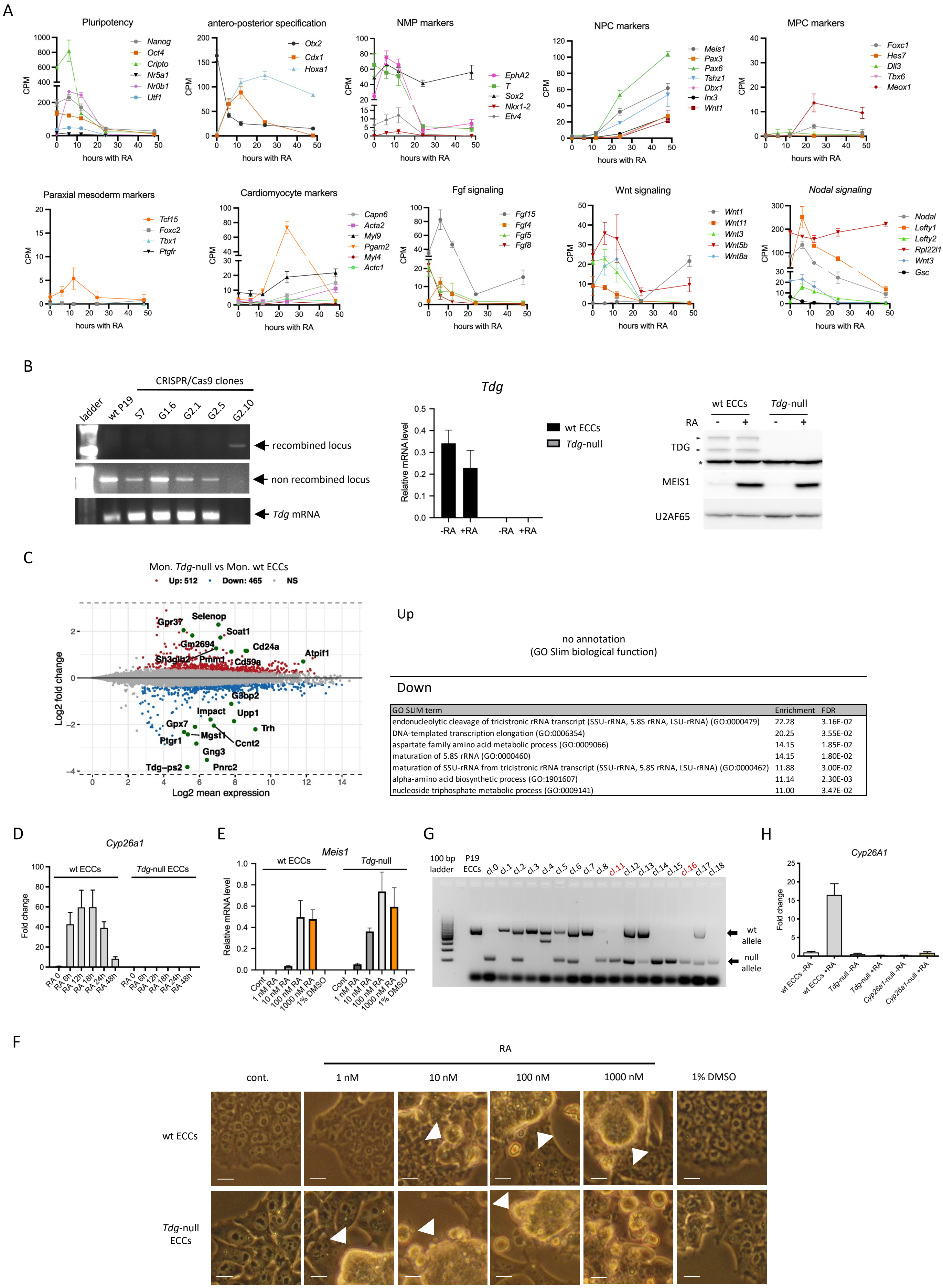

| GO SLIM term | Enrichment | FDR |
|---|---|---|
| endonucleolytic cleavage of tricistronic rRNA transcript (SSU-rRNA, 5.8S rRNA, LSU-rRNA) (GO:0000479) | 22.28 | 3.16E-02 |
| DNA-templated transcription elongation (GO:0006354) | 20.25 | 3.55E-02 |
| aspartate family amino acid metabolic process (GO:0009066) | 14.15 | 1.85E-02 |
| maturation of 5.8S rRNA (GO:0000460) | 14.15 | 1.80E-02 |
| maturation of SSU-rRNA from tricistronic rRNA transcript (SSU-rRNA, 5.8S rRNA, LSU-rRNA) (GO:0000462) | 11.88 | 3.00E-02 |
| alpha-amino acid biosynthetic process (GO:1901607) | 11.14 | 2.30E-03 |
| nucleoside triphosphate metabolic process (GO:0009141) | 11.00 | 3.47E-02 |

Suppl. Fig. 1

A

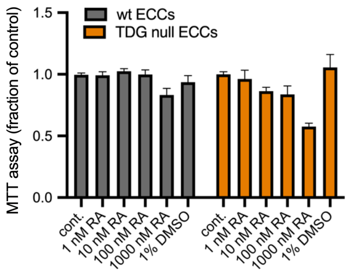

B

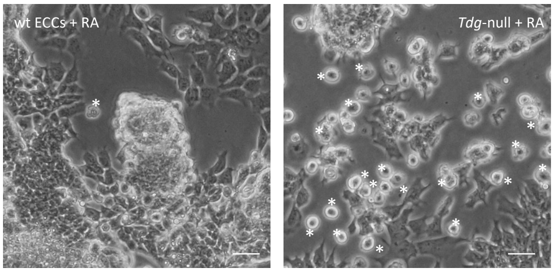

C

| Gene name | Effect on ferroptosis | Log2FC (mon. no RA) | Adj P value |
|---|---|---|---|
| *Cxcr2* | - | 0.74 | 4.23e-12 |
| *Emp2* | - | 2.03 | 1.11e-12 |
| *Eno1* | - | 0.71 | 8.10e-13 |
| *Gm2694* | + | 1.87 | 5.99e-17 |
| *Gpx7* | - | -2.16 | 1.44e-17 |
| *Meg3* | - | -2.96 | 1.05e-10 |
| *Mgst1* | - | -2.52 | 4.66e-16 |
| *Park7* | - | 0.70 | 4.15e-12 |
| *Phb2* | + | -0.74 | 6.60e-11 |
| *Ptgr1* | - | -2.42 | 2.55e-14 |
| *Selenop* | - | 2.32 | 1.43e-39 |
| *Soat1* | - | 1.76 | 2.02e-26 |
| *Stmn1* | - | 0.68 | 1.17e-09 |
| *Upp1* | + | -1.876 | 1.28e-43 |
| *Vamp8* | - | 0.87 | 8.97e-10 |

D

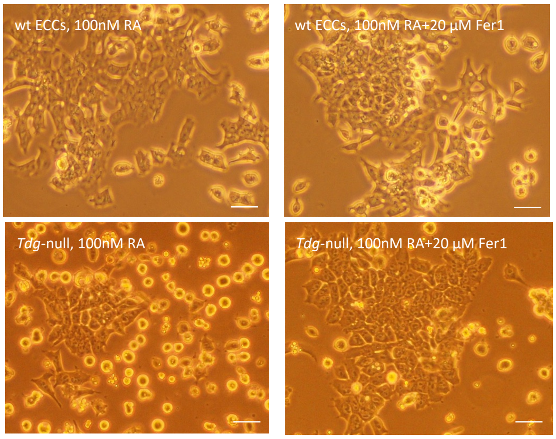

Suppl. Fig. 2

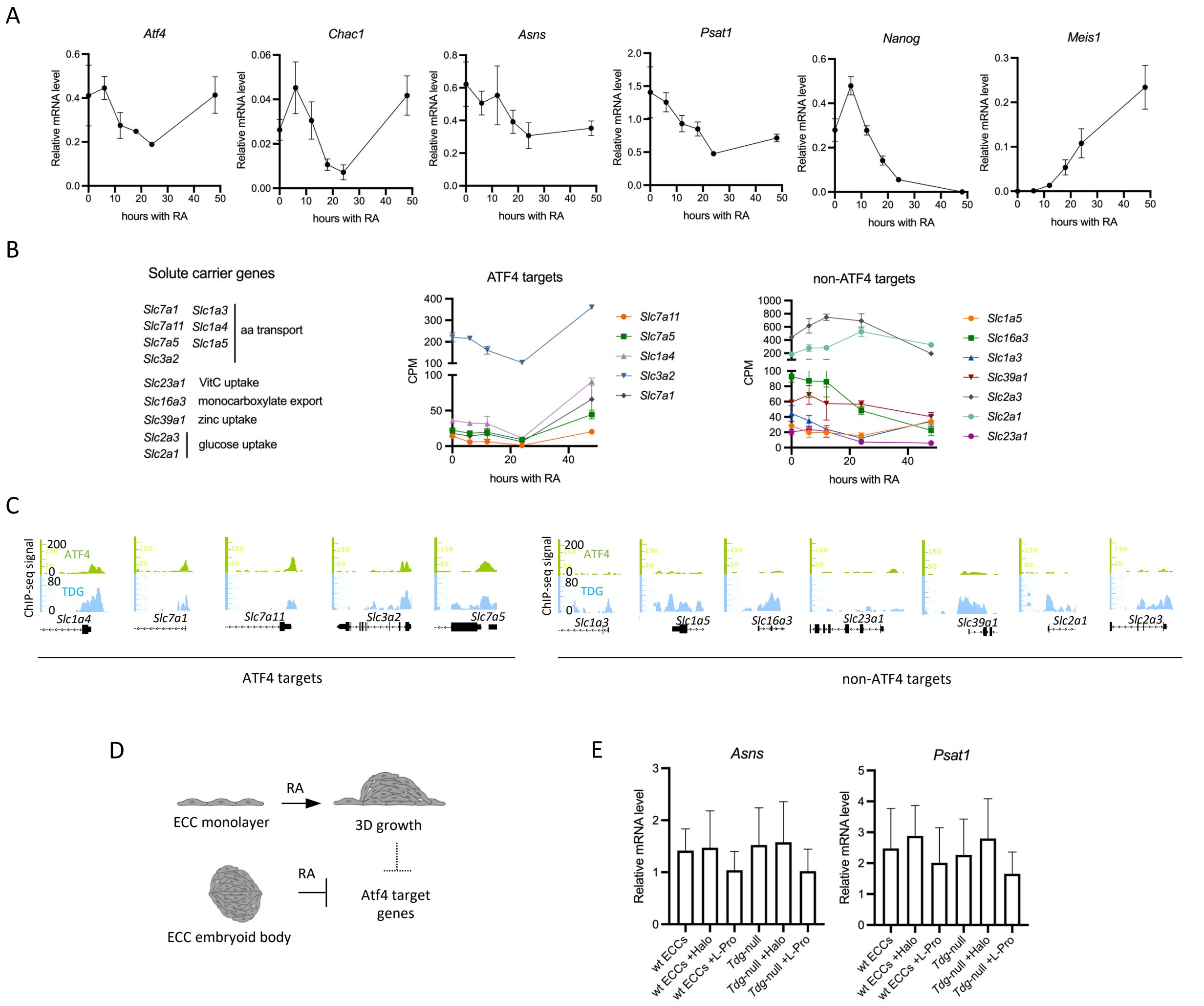

F

| | | *Tdg*-null vs wt ECCs | | | | Adh wt ECCs | | Adh *Tdg*-null | |
|---|---|---|---|---|---|---|---|---|---|
| | | EB no RA | | Adh no RA | | RA vs no RA | | RA vs no RA | |
| | Gene | Log2FC | Adj. Pval | Log2FC | Adj. Pval | Log2FC | Adj. Pval | Log2FC | Adj. Pval |
| Activation | *Mgst1* | -2.467 | 1.294e-8 | -2.524 | 4.665e-16 | 0.211 | 0.491 | -1.102 | 0.059 |
| | *Rhebl1* | -0.155 | 0.695 | -0.044 | 0.936 | -1.242 | 1.542e-11 | -1.500 | 3.596e-13 |
| | *Rheb* | -0.192 | 0.526 | -0.117 | 0.727 | 0.359 | 0.021 | 0.181 | 0.402 |
| | *Gpx7* | -2.215 | 6.522e-13 | -2.159 | 1.440e-17 | -0.483 | 0.041 | -0.941 | 0.023 |
| | *Rpl22l1* | -1.365 | 5.167e-17 | -1.108 | 1.4509e-12 | -0.160 | 0.416 | -1.213 | 7.802e-12 |
| | *Slc7a5* | -0.849 | 0.104 | 0.219 | 0.708 | 0.085 | 0.842 | -2.624 | 2.370e-14 |
| Inhibition | *G3bp2* | -1.503 | 8.067e-21 | -1.159 | 7.656e-17 | -0.400 | 0.004 | 0.217 | 0.280 |
| | *Tsc1* | -0.482 | 0.553 | -0.106 | 0.925 | 0.030 | 0.963 | -0.653 | 0.237 |
| | *Tsc2* | 0.207 | 0.737 | 0.190 | 0.778 | 0.081 | 0.860 | 0.208 | 0.633 |
| | *Sesn1* | 0.600 | 0.114 | 0.108 | 0.878 | -1.471 | 1.797e-6 | -0.060 | 0.899 |
| | *Deptor* | 0.110 | 0.902 | 0.131 | 0.883 | -1.888 | 3.377e-6 | -1.765 | 4.864e-5 |

Suppl. Fig. 3

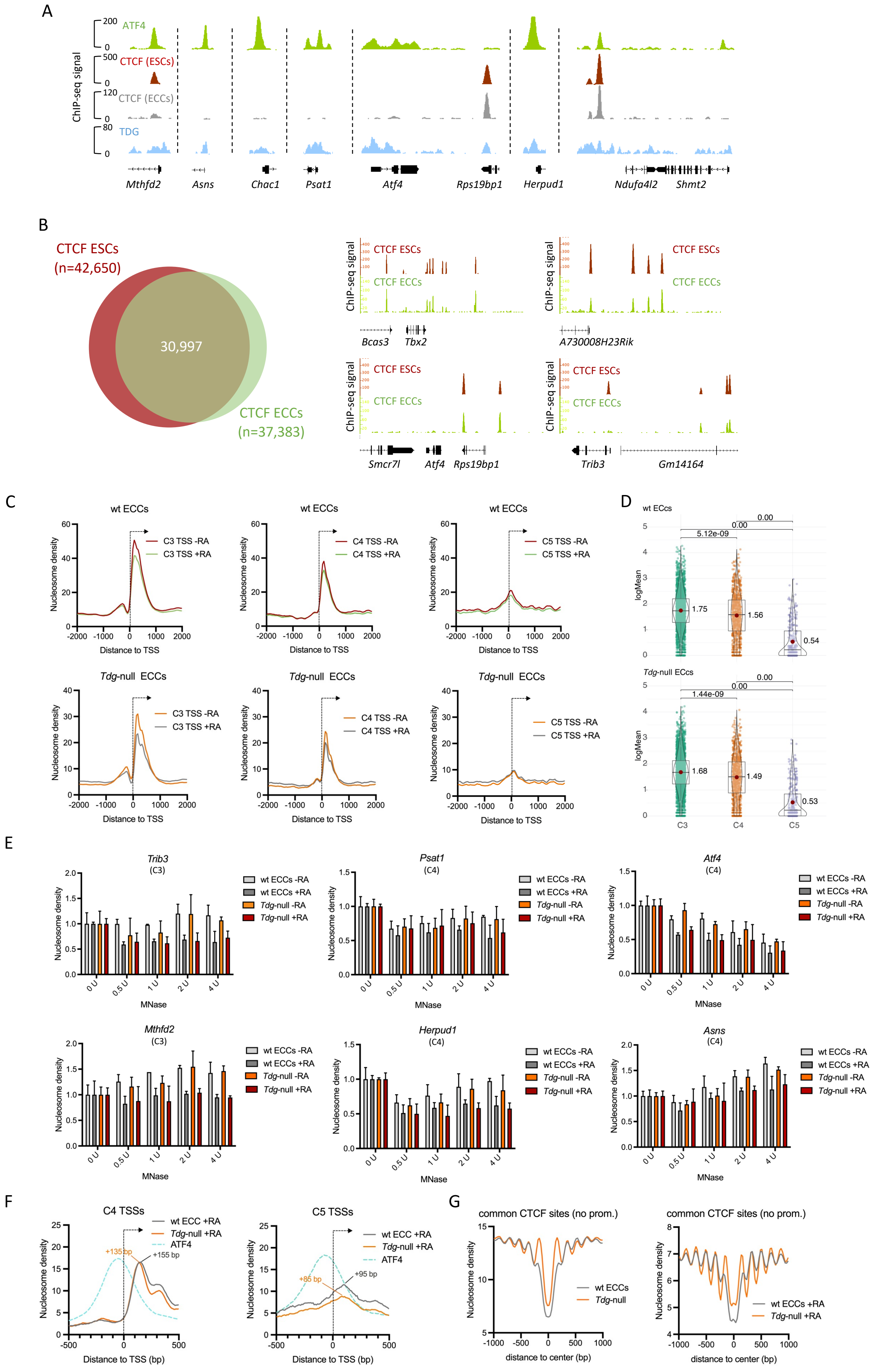
A
ChIP-seq signal
ATF4
CTCF (ESCs)
CTCF (ECCs)
TDG
Mthfd2
Asns
Chac1
Psat1
Atf4
Rps19bp1
Herpud1
Ndufa4l2
Shmt2
B
CTCF ESCs (n=42,650)
30,997
CTCF ECCs (n=37,383)
Bcas3
Tbx2
A730008H23Rik
Smcr7l
Atf4
Rps19bp1
Trib3
Gm14164
C
wt ECCs
Tdg-null ECCs
C3 TSS -RA
C3 TSS +RA
C4 TSS -RA
C4 TSS +RA
C5 TSS -RA
C5 TSS +RA
Nucleosome density
Distance to TSS
D
wt ECCs
Tdg-null ECCs
logMean
5.12e-09
1.44e-09
0.00
1.75
1.56
0.54
1.68
1.49
0.53
C3
C4
C5
E
Trib3 (C3)
Psat1 (C4)
Atf4 (C4)
Mthfd2 (C3)
Herpud1 (C4)
Asns (C4)
wt ECCs -RA
wt ECCs +RA
Tdg-null -RA
Tdg-null +RA
MNase
F
C4 TSSs
C5 TSSs
wt ECC +RA
Tdg-null +RA
ATF4
+135 bp
+155 bp
+85 bp
+95 bp
Distance to TSS (bp)
G
common CTCF sites (no prom.)
wt ECCs
Tdg-null
wt ECCs +RA
Tdg-null +RA
distance to center (bp)

Suppl. Fig. 4

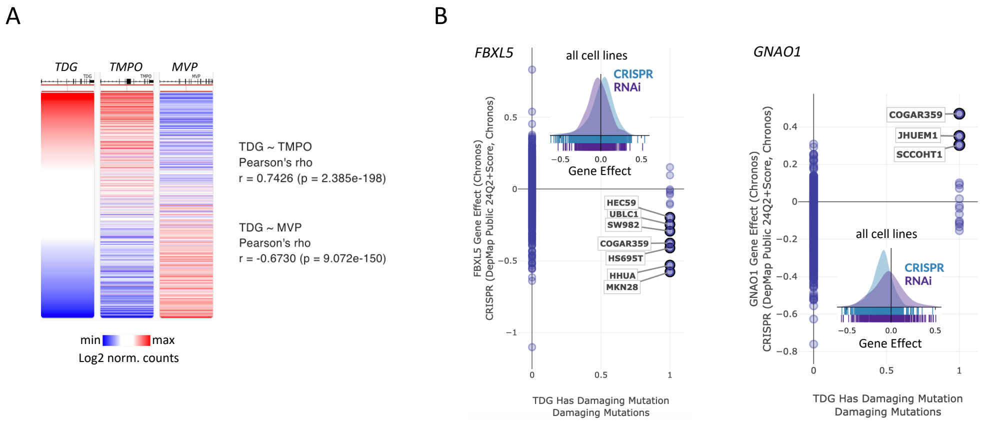
A
TDG
TMPO
MVP
TDG ~ TMPO
Pearson's rho
r = 0.7426 (p = 2.385e-198)
TDG ~ MVP
Pearson's rho
r = -0.6730 (p = 9.072e-150)
min
max
Log2 norm. counts
B
FBXL5
all cell lines
CRISPR
RNAi
Gene Effect
FBXL5 Gene Effect (Chronos)
CRISPR (DepMap Public 24Q2+Score, Chronos)
HEC59
UBLC1
SW982
COGAR359
HS695T
HHUA
MKN28
TDG Has Damaging Mutation
Damaging Mutations
GNAO1
COGAR359
JHUEM1
SCCOHT1
all cell lines
CRISPR
RNAi
Gene Effect
GNAO1 Gene Effect (Chronos)
CRISPR (DepMap Public 24Q2+Score, Chronos)
TDG Has Damaging Mutation
Damaging Mutations

Suppl. Fig. 5

**Supplementary Table 1: list of oligonucleotides used in this study**

| Primer type | Gene name | Forward primer | Reverse primer |
|---|---|---|---|
| | *RPS28* | GGTGACGTGCTCACCCTATT | TTCCGTGGGCTAAGTAGTGG |
| | *Irx3* | ACAGCACCTGCTGGGACTT | TTCTCCACTTCCAAGGCACT |
| | *Pax6* | ATGGCAAACAACCTGCCTAT | CATGGGCTGACTGTTCATGT |
| | *Dbx1* | CTGAAGTTTGGGGTGAATGCC | CGAACGTCTTGGGAGGAGG |
| | *Oct4* | TTGCAGCTCAGCCTTAAGAA | TGGTCTCCAGACTCCACCTC |
| | *Nanog* | AAGCAGAAGATGCGGACTGT | GCACTTCATCCTTTGGTTTTG |
| | *Meis1* | CAGAAAAAGCAGTTGGCACA | TGCTGACCGTCCATTACAAA |
| | *Wnt1* | GAACTGCCCCACTGCTCC | GCGGAGGTGATTGCGAAGAT |
| | *Wnt11* | ACAACAGTGAAGTGGGGAGAC | GTGCGGATGGAGCAGGAG |
| **RT-qPCR** | *Brachyury* (*T*) | GCTTTCCCGAGACCCAGTTC | GGCATCAAGGAAGGCTTTAGC |
| | *Cyp26a1* | CCCATTGACGTGCCCTTTAG | ATATCCAGCCTCTCTCCCCT |
| | *Tdg* | CCCCGATCCTGTGGTGAGTC | CCAGTCGTGACTGCCATGT |
| | *Acta2* | ATGTCCCCGCCATGTATGTG | CCATCTCCAGAGTCCAGCAC |
| | *Asns* | CGTATCCTTTTTGACAACGCTATCA | GCAGAGAGGCAGCAACCAAG |
| | *Atf4* | GCTATGGATGATGGCTTGGC | GCATCGAAGTCAAACTCTTTCAGA |
| | *Bcat1* | CTGCCAGCCTCTACATCCG | CGGTCTTCAAGGAGGGTCAC |
| | *Bdnf* | TACCTGGATGCCGCAAACAT | GCTGTGACCCACTCGCTAAT |
| | *Chac1* | GGGCTTCGTTCGTGGCTATA | GAGGCGGTAGAACTCAAAGA |
| | *Mthfd2* | TGCTTGGACCAGTACTCTATGC | CTGCCAGCCACTACCACAT |
| | *Psat1* | TTGGAACGGTGAACATTGTCC | GCGTCCGGGTTGAGGTTC |
| | *Nodal* | CCTCTGGCGTACATGTTGAG | GGTCACGTCCACATCTTGC |
| | *Lefty1* | GCTCACACAAGTTGGTTCGT | TGAGCTCCATAGTCCTTGAGG |
| | *Lefty2* | ACACAAGTTGGTCCGTTTCG | GAGCTCCGTAGTCCTTGAGG |
| | *Gpx4* | CTCATTGATAAGAACGGCTGCG | GATAGCACGGCAGGTCCTTC |
| | *Gpx7* | CCCTGCCTTCAAGTACCTAACC | CTCCCACCACCTTTCCGTC |
| | *Mgst1* | CCGCATTCCAGAGGATAACCA | TTCCACCTTCTCGTCAGTGC |
| | *Asns* | CAGAGAAGGTGGAGCAGAGG | ATGATGAAACTTCCCGCACG |
| | *Atf4* | CGCCTTGTAAGACACCGGAA | GCCCCATGAGAGAAAAGTGC |
| **ChIP-qPCR** | *Bcat1* | GGGTGCAAATGTGAGTCTCC | GCCCAGCTCTCCATCTTCC |
| | *Chac1* | CTGACGCAATCTGACTCGCC | CTCCTCCTCCCGTGGCAG |
| | *Mthfd2* | AAGGCTAGAACTGGTGGGCT | TACCAACTTCCCTCCTCCCG |
| | *Psat1* | ACGATCAATCAACTCCTCCTGG | GGTGGCTGGTGCAATCCC |
| | *Nampt*-nuc1 | CCCATTTTTCTCCTTGCTCGC | GATGTTGAACTCGGCTTCTGC |
| | *Nampt*-nuc2 | GACTGGATGAGACCGTGGGA | ACACCGCCAGGACCTCTC |
| | Negative ctrl | TCCTTTGACCTCCACACACA | GCAACAGAAGATGCGATGAA |

| | | | |
|---|---|---|---|
| | *Asns* | GACAGCACATCCTCCGGC | CTGCTCCACCTTCTCTGGC |
| | *Atf4* | GAAGTGTTGGCGGGGGAC | AAGTGCTTGGCCACCTCC |
| **MNase-qPCR** | *Herpud1* | CCGTTCGGCATCCCTGAG | GCGCTGATTGGGACTCTTCA |
| | *Mthfd2* | CACACTCACCGCACTGCC | TTCCTTGTTGTCTGCGTTGG |
| | *Psat1* | CACCTTACCGAGTGTGGCAG | GCTGTCGCCTTAGCACCAT |
| | *Trib3* | TACCTTGTCCTGGAGCCTCA | CACTAGCGTGCAGGAGACTC |
| | Ctrl | ATCTGGAAGCAGACGGACAC | TGGAGGCAAAAGAGAAGCTC |

| Primer type | Name | Sequence |
|---|---|---|
| | *Cyp26a1* gRNA105 sense | CACCGGAGGGCGCAGCTGCGATCG |
| | *Cyp26a1* gRNA105 antisense | AAACCGATCGCAGCTGCGCCCTCC |
| **CRISPR/Cas9** | *Cyp26a1* gRNA170 sense | CACCGCGCGTCGGTGCGCACCATCC |
| **Gene Inactivation** | *Cyp26a1* gRNA170 antisense | AAACGGATGGTGCGCACCGACGCGC |
| | *Cyp26a1*_screen_PCR_up | GCTCTGCACCTTCGTGCT |
| | *Cyp26a1*_screen_PCR_dw | GAATCGTGCAGGTTGGAGAG |
| | *Tdg* non-recombined_locus_up | AGCCCCACTTGGGTAAGACT |
| | *Tdg* non-recombined locus_dw | ACACTTACCCGTGCGTTAGC |
| | *Tdg* recombined locus_up | GCCTGAAGAACGAGATCAGC |
| | *Tdg* recombined locus_dw | CAAACCCCTCTGAAAGTTGG |
| | *Tdg* gRNA1 target sequence | GCGCCCGCGTTACCTGCGCG |
| | *Tdg* gRNA2 target sequence | CTCGCGGCTGGGGTCGGTCA |
